# Using pseudoalignment and base quality to accurately quantify microbial community composition

**DOI:** 10.1101/097949

**Authors:** M. Reppell, J. Novembre

## Abstract

Pooled DNA from multiple unknown organisms arises in a variety of contexts, for example microbial samples from ecological or human health research. Determining the composition of pooled samples can be difficult, especially at the scale of modern sequencing data and reference databases. Here we propose the novel pooled DNA classification method Karp. Karp combines the speed and low-memory requirements of k-mer based pseudoalignment with a likelihood framework that uses base quality information to better resolve multiply mapped reads. In this text we apply Karp to the problem of classifying 16S rRNA reads, commonly used in microbiome research. Using simulations, we show Karp is accurate across a variety of read lengths and when samples contain reads originating from organisms absent from the reference. We also assess performance in real 16S data, and show that relative to other widely used classification methods Karp can reveal stronger statistical association signals and should empower future discoveries.

## 1. Introduction

The study of microbial community composition has been revolutionized by modern genetic sequencing. Experimenters can forgo the laborious work of culturing cells and detect a broader range of taxa than was previously possible. This improved ability to describe the microbes present in a pooled sample has led to important findings in human health (Davenport *et al*., 2014; Wu *et al*., 2011; Turnbaugh *et al*., 2009) and ecology (Metcalf *et al*., 2016; Godon *et al*., 2016). These findings rely on quantification of the taxa present in experimental samples, and towards that goal many methods have been developed. The ever-increasing scale of both sequencing data and relevant reference databases require that such methods be efficient in addition to accurate. Here we present a novel method, Karp, which combines the speed of k-mer-based pseudoaligning with a likelihood framework that incorporates base quality information. In this work we use Karp to classify the taxonomy of pooled 16S microbiome data quickly and with an accuracy superior to widely adopted alternative methods.

Microbiome samples are commonly generated using either shotgun sequencing or the sequencing of marker genes, most often the gene encoding 16S ribosomal RNA. Classifying the output of shotgun sequencing can be difficult, as limited reference databases exist for entire bacterial genomes, so whole genome sequencing generally either requires computationally intensive de novo assembly methods (Cleary *et al*., 2015; Howe *et al*., 2014; Boisvert *et al*., 2012) or limits the range of organisms available for study (Scholz *et al*., 2016). Alternatively, several large reference databases exist for microbial 16S sequences (Cole *et al*., 2014; Quast *et al*., 2013; DeSantis *et al*., 2006). The 16S gene contains alternating regions of highly conserved and highly variable sequences, making it easy to target and well powered for differentiating taxa. Many experiments target one or several of the 16S hypervariable regions and sequence to a high depth (Howe *et al*., 2016; Ahn *et al*., 2011; Chakravorty *et al*., 2007).

Sequence identification problems can be broadly classified as either open-reference or closed-reference. In open-reference problems the sequences of possible contributors are unknown. In a closed-reference problem the sequences of contributors are known, and classification is typically a process of matching the observed sequencing reads against a reference database. Closed-reference methods for classifying microbial samples face several significant challenges. First, methods must be able to provide unbiased estimates when samples contain previously unidentified taxa. Second, microbial samples often contain a range of genetic diversity unmatched by single species sequencing samples. And finally, methods must efficiently compare sequences against reference databases containing potentially millions of organisms.

Microbiome classification tools can generally be divided into three categories. The first is based on similarity scores between a query and potential references. Many early similarity based methods first employed the Basic Local Alignment Search Tool (BLAST) (Altschul *et al*., 1990), which calculates both a similarity score and relative significance for local alignments of queries against reference sequences. Several methods refined BLAST output to classify sequence origin (Glass *et al*., 2010; Horton *et al*., 2010; Huson *et al*., 2007), however, the BLAST algorithm is computationally very intensive, making methods based on it hard to scale with both reference panel size and sequencing depth. These early BLAST based methods have largely been superseded by the USEARCH and UCLUST algorithms (Edgar, 2010, 2013) and several other recent similarity-based clustering algorithms (Al-Ghalith *et al*., 2016; Albanese *et al*., 2015; Mahe *et al*., 2014; Kopylova *et al*., 2012) that are fast enough to handle modern data (millions of reads, each one hundreds of base pairs long). The speed and accuracy of these modern clustering algorithms has been shown to be very similar (Kopylova *et al*., 2016; Al-Ghalith *et al*., 2016). A second approach for classifying sequences is based on the shared phylogeny of samples, and places query sequences along a phylogenetic tree. Phylogenetic methods using maximum-likelihood estimation (Berger *et al*., 2011), Bayesian posterior probabilities (Matsen *et al*., 2010), or neighbor-joining (Price *et al*., 2009) have all been developed. While representing the explicit relationships between organisms provided by phylogenetic methods is attractive, these methods impose a large computational burden. Also, while they often make accurate taxonomic assignments, phylogenetic methods tend to suffer from low sensitivity (Bazinet and Cummings, 2012). The third category consists of methods that use sequence composition to classify. Early sequence composition methods calculated the probability of a query originating from a specific taxon based on shared k-mers (Rosen *et al*., 2008; McHardy *et al*., 2007; Wang *et al*., 2007). In a review of early methods Bazinet *et al*. (2012) found that the sequence composition method Naive Bayes Classifier (NBC)(Rosen *et al*., 2008), had the best balance of sensitivity and specificity; but NBC is too slow for large reference databases. Recently the Mothur pipeline (Kozich *et al*., 2013) provides an implementation of the k-mer based Wang *et al*. (2007) naive Bayes algorithm that can be effectively run on large numbers of reference and query sequences. Kraken (Wood and Salzberg, 2014) and CLARK (Ounit *et al*., 2015) are two additional recent sequence-composition methods designed for modern datasets that had the highest accuracy in a third party evaluation (Lindgreen *et al*., 2016). However, both Kraken and CLARK require powerful workstations (>75 GB RAM) with substantial hard-drive space to run in their most accurate modes.

Very recently, the development of pseudoalignment (Bray *et al*., 2016) has allowed sequence composition classification with minimal computational requirements and an accuracy superior to both Kraken and CLARK (Schaeffer *et al*., 2015; Teo and Neretti, 2016). Pseudoaligning, originally developed in the context of RNA sequencing experiments, is a rapid k-mer based classification that uses a de Bruijn Graph of the reference database to identify potential matches for a query sequence without aligning the query to reference sequences. Pseudoaligning is very fast, and is implemented in the software Kallisto (Bray *et al*., 2016), which uses an expectation maximization (EM) algorithm to resolve multiply-mapped reads without assigning them to a single taxonomic unit. The speed advantages of Kallisto and pseudoaligning come at a cost; notably it ignores information about sequencing quality that could help assign multiply-mapped reads more accurately. Sequencing errors occur non-uniformly along reads, and base-quality scores record the probability of errors at each base. Thus, classification can be improved by using base-quality scores to help distinguish true mismatches between reads and references from sequencing errors.

Kallisto’s limitations led us to develop Karp, a program that leverages the speed and low memory requirements of pseudoaligning with an EM algorithm that uses sequencing base-quality scores to quickly and accurately classify the taxonomy of pooled microbiome samples. Here, we demonstrate with simulations of 16S sequencing experiments the improvement in accuracy that Karp provides relative to Kallisto, as well as modern similarity-based methods (using Quantitative Insights Into Microbial Ecology (QIIME)), and the Wang *et al*. (2007) naive Bayesian classifier (using Mothur). We also use simulations to demonstrate how Karp leads to better estimates of important summary statistics and remains robust when sequences from organisms absent from our reference database are present at high frequencies in samples. Finally, we assess performance in a real 16S dataset with 368 samples drawn from two individuals over two days. In this data Karp finds more taxa with stronger association signals that differ between the two individuals. Karp also maintains comparable classification errors when a random forest is employed to classify which location or individual each sample originated from.

## 2. Methods

### 2.1 An overview of Karp

The aim of Karp is to estimate a vector 𝓕 = (*f*_1_, …*f_M_*), containing the proportion of a pooled DNA sample that is contributed by each of *M* possible reference haplotypes. Figure 1 gives an outline of Karp’s classification process. The first step in using Karp is the construction of a k-mer index of the *M* reference sequences. This index catalogs the subset of the *M* reference haplotypes that contain each unique k-mer of a given length. Next, the query reads are pseudoaligned using the k-mer index. Query reads that pseudoalign to multiple references (multiply-mapped reads) are locally aligned to each potential reference, and each reference’s best alignment is kept. Queries that pseudoalign to a single reference are assigned without alignment. Next, for multiply-mapped reads the likelihood that they originated from each potential reference is calculated using the best alignment and the base-quality scores that correspond to the read. After the likelihoods for every query read have been calculated, an EM-algorithm is used to estimate the relative frequencies of each reference haplotype contributing to the pool. More details about the method are provided in the following sections.

**Figure 1:**
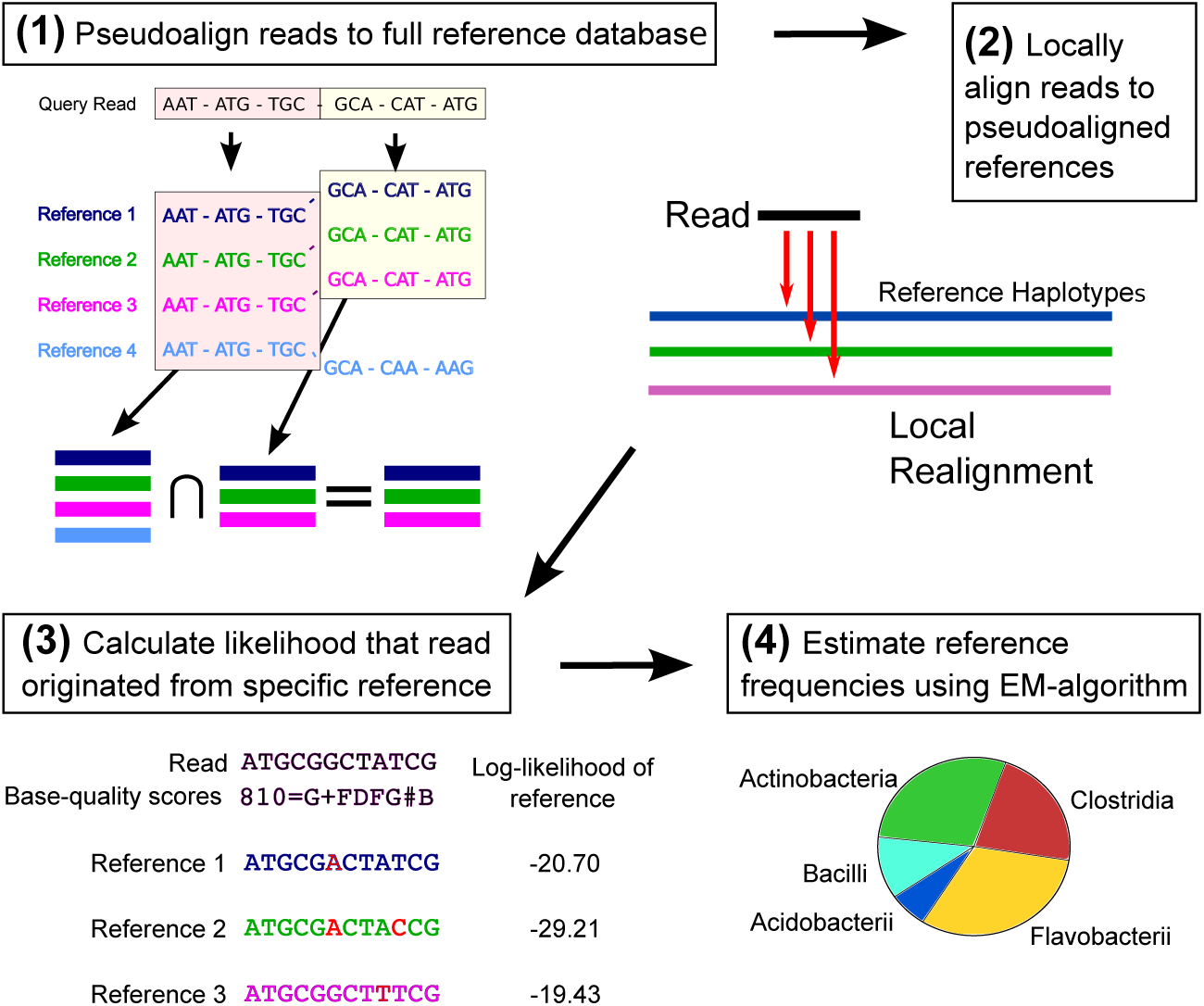
An overview of Karp. (1) Query reads are pseudoaligned against an index of the reference database, resulting in a set of references they could have potentially originated from. (2) The query reads are locally aligned to the possible references. (3) Using the best alignment, the likelihood that a read originated from a specific reference is calculated. (4) Using the read likelihoods an EM-algorithm is employed to estimate the relative abundances of the reference haplotypes in the pool of query reads.

### 2.2 Pseudoaligning and alignment

Aligning millions of reads against hundreds of thousands of references is impractical in both memory and time. However, calculating the probability that a read originated from a given reference sequence using base-quality information requires an alignment. To overcome this challenge, Karp uses pseudoalignment as a filter before performing local alignment. Pseudoalignment is a fast and memory efficient way to narrow the space of possible references from which a query read may have originated. Our pseudoaligning algorithm is directly based on that of Kallisto (Bray *et al*., 2016). Briefly, first an indexed de Bruijn Graph of the reference database is constructed, with each observed k-mer mapped to an equivalence class of reference sequences that it is contained in. Next, each query read is decomposed into its constituent k-mers, which are searched against the index. An intelligent coding of the index allows for a minimal number of k-mer look-ups. Kallisto uses a strict intersection of the equivalence classes returned by the k-mer search to arrive at a pseudoalignment. Karp can also be set to use the strict intersection of equivalence classes. However, because mismatched bases are accounted for in Karp’s read likelihood framework, we are more concerned with false negatives than false positive matches and the default setting is more inclusive. In Karp’s default mode, if no strict intersection is observed, the intersection of all equivalence classes with the same maximum number of matched k-mers, conditional on the maximum being > 1, are declared matches. Reads with < 2 matched k-mers are always removed from analysis for failing to pseudoalign.

After pseudoaligning, reads are locally aligned to the matching reference sequences using the Striped Smith-Waterman algorithm (Zhao *et al*., 2013; Farrar, 2007) (SSW penalties: mismatch (2), gap opening (3), gap extending (1)).

### 2.3 Read likelihoods

Our likelihood and EM frameworks build closely on the work of Kessner *et al*. (2013), whose software Harp implemented a method for estimating haplo-type frequencies in pooled DNA. Kessner *et al*. (2013) recognized the potential of their method to improve accuracy in microbiome studies, but Harp was computationally infeasible with modern reference databases.

For a read *r_j_* with *j* ∈ 1, …, *N* and length *L_j_*, let (*r_j_* [1], …, *r_j_* [*L_j_*]) be the base calls at each position along the read. Assume we have a reference database with *M* possible haploid reference sequences, which we will refer to as reference haplotypes. For reference haplotype sequence *h_k_* with *k* ∈ 1,…,*M* let (*h*_*k*,*j*_[1],…, *h*_*k*,*j*_[*L_j_*]) give the values of the bases in *h_k_* corresponding to the best alignment of read *r_j_*. Note the entries in this vector may not be contiguous due to insertions, deletions, or because the reads are paired-end. Define the probability of sequencing error at each position as *q_j_* [*i*] = *P*(*r_j_* [*i*] ≠ *h*_*k*,*j*_ [*i*]) for *i* ∈ 1, …, *L_j_*, and define the variable *η_j_*, a vector of length *M* with components *η*_*j*,*k*_ = 1 if *r_j_* originated from haplotype *h_k_* and 0 otherwise. Assuming sequencing errors are independent, we can then formulate the probability of read *r_j_* arising from reference *h_k_*, which we label *l*_*j*,*k*_, as

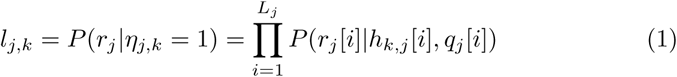

where, if we assume every base is equally likely when an error occurs

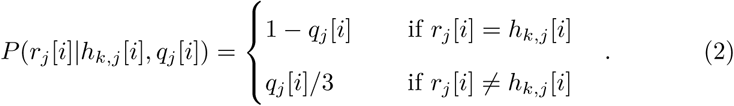

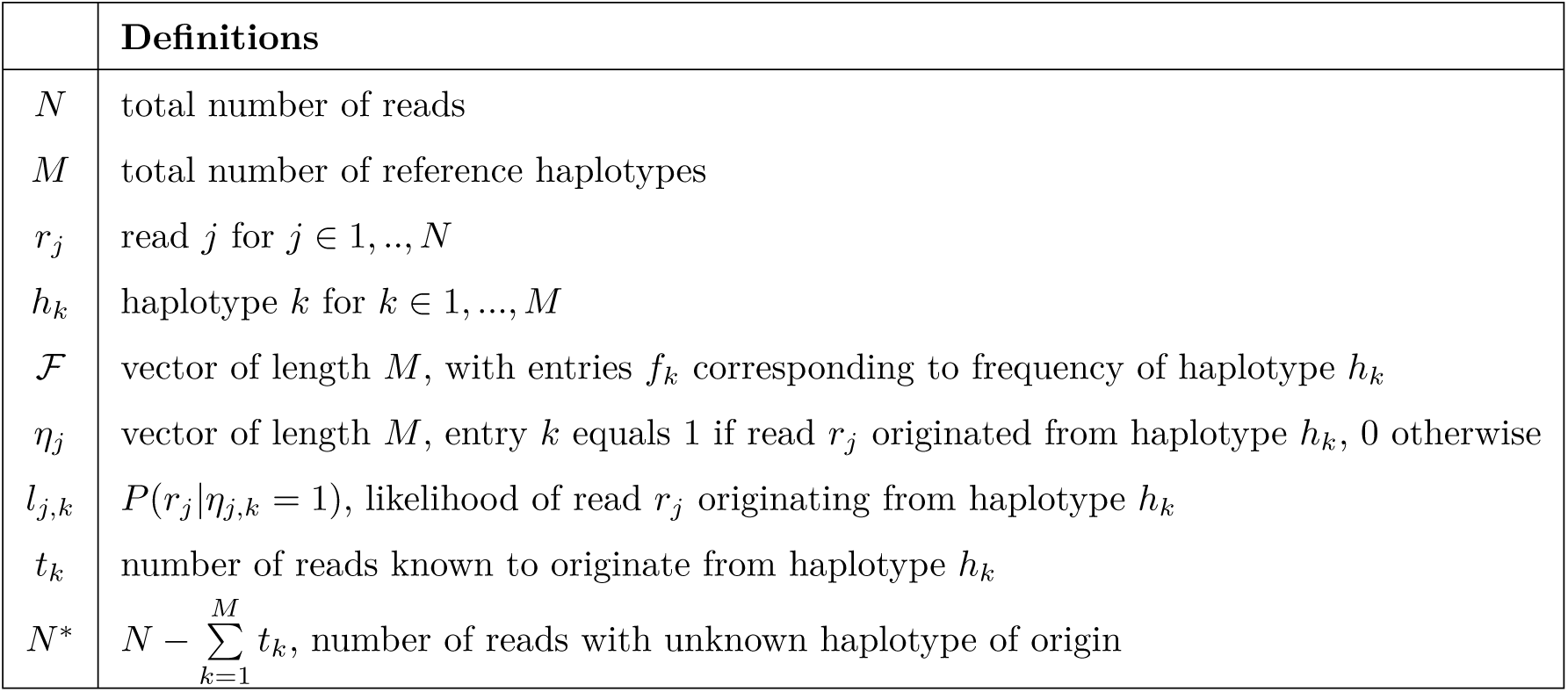

A more complete definition of the probability would sum over all possible alignments of *r_j_* to *h_k_*. However, in non-repetitive marker gene sequence the best local alignment typically contributes such a large proportion of the probability weight, that excluding alternate local alignments has a negligible impact on results but substantially improves computation.

### 2.4 Estimating reference haplotype proportions

As previously noted, the aim of our method is to estimate a vector 𝓕 = (*f*_1_, …,*f_M_*), containing the frequencies of the *M* possible reference haplotypes in a pooled DNA sample. If we were to observe which reference haplotype gave rise to each read in our sample, the maximum likelihood estimate of 𝓕, 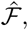 would follow directly from the multinomial likelihood. In reality, we observe the reads *r*, but the reference haplotypes that they originate from, *η*, are unobserved. To estimate 𝓕 we therefore employ an EM algorithm, with a form common to mixture model problems. Details of our EM algorithm are provided in supplementary section 7.1.

Karp modifies the standard mixture EM algorithm in two ways to speed up performance. The first is an assumption that if a read *r_j_* uniquely pseudoaligns to a reference *h_k_* then *P*(*η*_*j*,*k*_ = 1|*r*, 𝓕) = 1. For haplotype *h_k_*, label the number of reads that uniquely map as *t_k_* and define 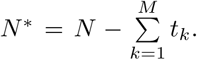 Then we can write the likelihood of the data as

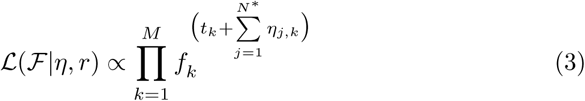

and our update step as

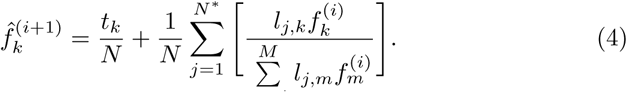

This assumption also provides a logical initial estimate of 𝓕^(0)^

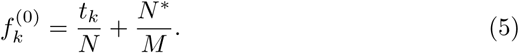

The second speed-up that Karp uses is an implementation of SQUAREM (Varadhan and Roland, 2004), which accelerates the convergence of EM algorithms by using information from multiple previous parameter updates to improve the current EM update step.

Additionally, Karp allows the user to specify a minimum reference haplotype frequency. After the frequency of a reference falls below this threshold during the EM updates, its value is set to zero and its frequency weight is distributed evenly across the remaining references. While this step technically violates the guarantee of the EM algorithm to reach a local maximum of the likelihood function, in practice we find that when there is sufficient information to distinguish closely related species this approach imposes a sparsity condition which is effective for avoiding the estimation of spurious references at very low frequencies. When only limited information to distinguish between closely related species exists, for example in data generated from a single 16S hypervariable region, it can be better to set the minimum frequency very low to avoid eliminating true low frequency taxa with probability weights distributed evenly across indistinguishable OTUs. Supplementary figures S2, S3, and S4 explore the impact of different thresholds on the simulated and real data presented in this study.

### 2.5 Read likelihood filter

Our EM method relies on the fact that all the reads in our sample originated from haplotypes present in our reference database. In real data this assumption can be problematic; the classification of microbial taxonomy is an ongoing project and many taxons have yet to be identified or referenced. To preserve the accuracy of our frequency estimates in the presence of reads from haplo-types absent from our references we implemented a filter on the maximum read likelihood value (Kessner *et al*., 2013).

Specifically, using the base-quality scores of the query reads we calculate a “null” distribution of likelihood values corresponding to what we would observe if every query were matched to its true originating reference and every mismatched base was the result of sequencing errors. Then, after the local realignment step we filter out query reads where the greatest observed likelihood falls too far outside this distribution, as these are unlikely to truly match any of the reference sequences present in the database. Karp includes the option to output the maximum likelihood for each read, which can be used to determine the appropriate cutoff value. In our simulations, where a variety of empirical quality score distributions were encountered, cutoff values between −3.0 and −1.5 yielded similar results, a finding in line with Kessner *et al*. (2013), and which supports a default value of −2.0. In the real 16S data from Lax *et al*. (2015) we explored thresholds between −0.5 and −7.0, and generally those > −1.5 yielded the lowest classification error rates (Supplementary Figure S4 and Table 3). For more details about the filter see supplement 7.2.

### 2.6 Karp collapse mode

The default approach in Karp estimates the relative frequencies of the individual haplotypes present in the reference database. In many microbiome databases there is not a one-to-one relationship between reference haplotypes and taxonomic labels; multiple haplotypes share a single label. When little information exists to distinguish closely related haplotypes apart, estimating the relative frequencies at the taxon level rather than haplotypes can improve accuracy. To accommodate this, Karp includes a collapse option, which adds a step to the estimation procedure. When the collapse option is used, after pseudoalignment and local alignment Karp calculates the average likelihood for each taxonomic label, and uses these likelihoods in the EM algorithm to estimate taxonomic frequencies. This can be interpreted in a Bayesian context as the likelihood a read is from a taxon under a uniform prior of its true reference sequence within that taxa. Karp output in collapse mode provides counts at each taxonomic level from species to phylum. Because it is estimating the frequencies of fewer categories, collapse mode is often faster than Karp’s default.

### 2.7 Simulating 16S reads

To compare Karp with alternative methods we simulated pooled sequence samples. We used GreenGenes version 13.8 (DeSantis *et al*., 2006) as our reference database. The general simulation procedure was as follows. First, a fixed number of reference sequences were selected at random from Greengenes and a vector of frequencies corresponding to these references was generated by drawing from a Dirichlet distribution. Next, a predetermined number of reads were simulated. For each read a reference haplotype was drawn at random according to its frequency in the original frequency vector. Then, along the chosen reference sequence a read start position was selected uniformly and a number of bases corresponding to the desired read length were copied from the reference. In the case of paired-end reads, the distance between pairs was drawn as an upper-bound Poisson random variable with an empirically derived mean. Bases which would cause the read to extend past the end of the reference were excluded from being initiation points. Once a read’s bases were copied, a corresponding base-quality score vector was generated based on an empirical distribution of quality scores. To simulate 75bp single-end reads we used the publicly available Illumina-sequenced mock-community dataset from the Human Microbiome Project (Peterson *et al*., 2009). For simulating 151bp paired-end reads we used the quality scores observed in Illumina-sequenced microbiome samples collected from Amish and Hutterite mattresses (Stein *et al*., 2016). Finally, 301bp paired-end reads were simulated using scores from a sample of human saliva downloaded from Illumina’s BaseSpace platform (https://basespace.illumina.com/projects/17438426). Finally, errors were simulated along the read with probabilities corresponding to the base-quality score at each position and assuming that the three alternative bases were equally likely. After adding errors the read was added to the pooled sample, and the algorithm proceeded to the next read.

### 2.8 Simulations

In our simulations we used samples containing 1 × 10^6^ sequencing reads, a depth inspired by recent high-depth studies (Stein *et al*., 2016) and designed to demonstrate the computational feasibility of Karp. For each sample we selected 1,000 reference haplotypes randomly from GreenGenes and simulated reads following the approach of section 2.7. The Dirichlet distribution used to generate the sample frequency vectors had identical alpha values varied between 0.002 and 7. These parameter settings created samples with a broad range of Shannon Diversity values (Supplementary Figure S5). We simulated 110 samples with 75bp single-end reads, 130 samples with 151bp paired-end reads, and 170 samples with 301bp paired-end reads, each with a unique mix of 1,000 reference haplotypes from GreenGenes. With Kallisto and Karp the raw forward and reverse reads were directly classified. For the QIIME algorithms the script *join_paired_ends.py* was run and the resulting contigs were classified.

We simulated an additional 100 samples with 75bp single-end reads to compare how each method’s frequency estimates impacted the estimation of common sample summary statistics. Many statistics, such as *β* Diversity, summarize the sharing of taxa between samples, so instead of 1,000 unique taxa in each sample, we used a shared pool of 1,000 taxa for all 100 samples, and further increased the similarity between samples by introducing correlation between the reference frequencies. The reference haplotype frequencies for each sample were a linear combination of a random Dirichlet variable generated in a manner identical to the simulations above and the reference frequencies of the preceding sample. In this way the samples again covered the full range of Shannon Diversity values, however the frequencies of shared taxa was potentially much higher, providing a broader range of summary statistic values in the simulations.

Next we compared how the methods performed when the simulated samples contained reads generated from taxa that were absent from the reference database being used for classification. We selected one phylum (Acidobacteria), one order (Pseudomonadales), and one genus (Clostridiisalibacter) at random from the taxa in GreenGenes with more than 30 reference sequences. Then, for each missing taxa, we simulated 10 samples where 50% of the reads originated from 3 different members and at least 5% of the reads came from closely related taxa (kingdom Bactera for Acidobacteria, class Gammaproteobacteria for Pseudomonadales, and family Clostridiaceae for Clostridiisalibacter). Next, we create 3 reduced GreeGenes reference databases, each with one of the missing taxa (including all lower ranking members) expunged. Finally, we classified the simulated samples using both the appropriate reduced reference database and the full GreenGenes database.

Finally, we examined how sensitive our results were to the assumption that base-quality scores are accurate representations of the probability of sequencing error. Karp assumes that base-quality scores follow the Phred scale, where *Q* the probability of a sequencing error is 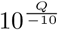 for quality score *Q*. Given that quality scores often overestimate the rate of errors, we simulated and classified 50 samples where the actual probability of an error was 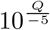 and also 50 samples where errors occurred uniformly at 1% of bases.

We compare the different classification methods using an AVGRE (AVerage Relative Error) metric (Schaeffer *et al*., 2015; Li, 2015; Sohn *et al*., 2014) which is based on the absolute value of the difference between the true and estimated counts of reads in the simulated samples. Define *M_a_* as the set of actual reference haplotypes contributing to a pooled sample and *M_e_* as the set of additional references a method classifies as having a non-zero number of reads that are not truly present. Also, let *T*_*i,e*_ be the count of reads estimated for reference *i* and *T*_*i,a*_ is the actual number of simulated reads from reference *i* present in the sample. Using these values the AVGRE metric has the form:

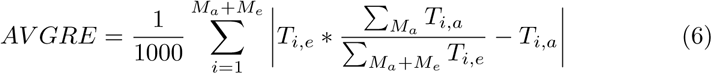

We include the scaling factor of 1/1000 in order to transform the value into an estimate of the average per-reference error rate, as our pooled simulation samples include 1,000 individual reference sequences. We use the same scaling factor when looking at errors in the estimation of higher order taxonomy for consistency, although the true number of references at any given taxonomic level will be < 1, 000.

### 2.9 Real data

To test the performance of Karp with real data we reanalyzed samples originally published by Lax *et al*. (2015). In brief, these samples were collected from the floor, shoes, and phones of two study participants every hour for two 12-hour time periods over the course of two successive days. From these samples the V4 region of the 16S rRNA gene was amplified and sequenced using the Illumina HiSeq2000 (llumina, San Diego, USA). Because this dataset contains many samples of known origin it is useful for assessing performance by measuring classification accuracy and the power to detect differences.

The data is publicly available at https://figshare.com/articles/\%/20Forensic_analysis_of_the_microbiome_of_phones_and_\%20shoes/1311743, and after download we used the scripts *split_libraries_fastq.py* and *extract_seqs_by_sample_id.py* from the QIIME software pipeline to demultiplex and split it into individual samples. After demultiplexing there were a total of 368 samples comprised of 151bp reads, with a median depth of 131,200 reads.

We classified the 368 samples using Karp, Kallisto, and UCLUST and performed three analyses. For each analysis, we used samples with a standardized depth of 25,000 reads, generated by subsampling without replacement. Five samples had < 25, 000 reads successfully classified by all three methods, leaving 363 samples for analysis: 103 phone samples, 207 shoe samples, and 53 floor samples. For our first analysis we used the randomForest package (Liaw and Wiener, 2002) in the program R to perform a random forest classification of the data using 1,000 trees and 6 different outcomes. We classified all the phone samples as coming from Person 1 or Person 2, did the same for all the shoe samples, then classified whether the phone samples from each person came from the front or back of their phones, and finally classified which of the shoe surfaces (front right, back right, front left, back left) each person’s shoe samples came from. We performed the subsampling and random forest classification 10 times, and calculated the average classification error for each analysis.

Next, we again subsampled 25,000 reads for each sample 10 separate times, and then performed a principal components decomposition (PCA) of the resulting matrices using the prcomp function in the R stats library. During the experiment, floor samples were collected alongside the shoe samples at the each time point. With the PCA decomposition we calculated the correlation between PCA 1 for the floor and shoe samples taken at the same time. For each method we calculated the average correlation across the 10 different subsampled matrices.

Finally, we tested for differences in the mean abundance of taxa between person 1 and person 2, first in the phone samples, and then between the shoe samples. After subsampling 25,000 reads for each sample, we tested each taxon with > 250 total reads across all samples using Welch’s t-test in R, and recorded the corresponding p-value and t-statistic. We used the p.adjust function in R to calculate false discovery rates (FDR) from the t-statistic p-values once all taxa had been tested.

### 2.10 Implementation

The program Karp is implemented in C++, and available for download from GitHub at https://github.com/mreppell/Karp. Karp takes as input sample fastq files, reference sequences in fasta format, and taxonomy files with labels corresponding to the references. The first stage of analysis with Karp is building a k-mer index of the references. Karp then uses this index along with the reference sequences to pseudoalign, locally align, and then quantify the taxonomy in the fastq file of query reads. Karp includes a post analysis option to tabulate multiple samples and calculate compositional summaries. Karp can make use of multi-threading to improve performance, and allows users to specify frequency thresholds, EM convergence conditions, likelihood filter parameters, and pseudoalignment k-mer length.

The simreads program we used to simulate sequence data with an empirical distribution of base-quality scores is available at https://bitbucket.org/dkessner/harp.

## 3 Results

### 3.1 Comparison of competing methods with simulations

To test the performance of Karp against alternatives we simulated 110 independent samples, each with 1 × 10^6^ 75bp single-end reads drawn from 1,000 reference haplotypes selected at random from the GreenGenes database. Each simulation used a unique set of 1,000 references, and the frequencies of each reference was varied to create a range of Shannon Diversity in the 110 samples (Supplemental Figure S5). We classified sequences against the full GreenGenes database using Karp, Kallisto, the Wang *et al*… (2007) Naive Bayes classifier implemented in Mothur, and several algorithms from QIIME including UCLUST, USEARCH, and SortMeRNA. We estimated errors as described in section 2.8.

At the level of individual reference haplotypes, which here we also refer to as operating taxonomic units or OTUs, we calculated estimation error for references with > 1 read present or classified. On average Karp had the lowest errors (34% smaller than Kallisto, 65 – 66% smaller than UCLUST, USEARCH, and SortMeRNA, Figure 2A). When we limited our comparison to references with a frequency > 0.1%, Karp remained the most accurate (errors 31% smaller than Kallisto, and 68–70% smaller than the QIIME algorithms, Figure 2B). The accuracy of all methods improved with increasing diversity, and Karp’s average error was 48% smaller when diversity was > 6.2 than when it was < 0.7.

**Figure 2:**
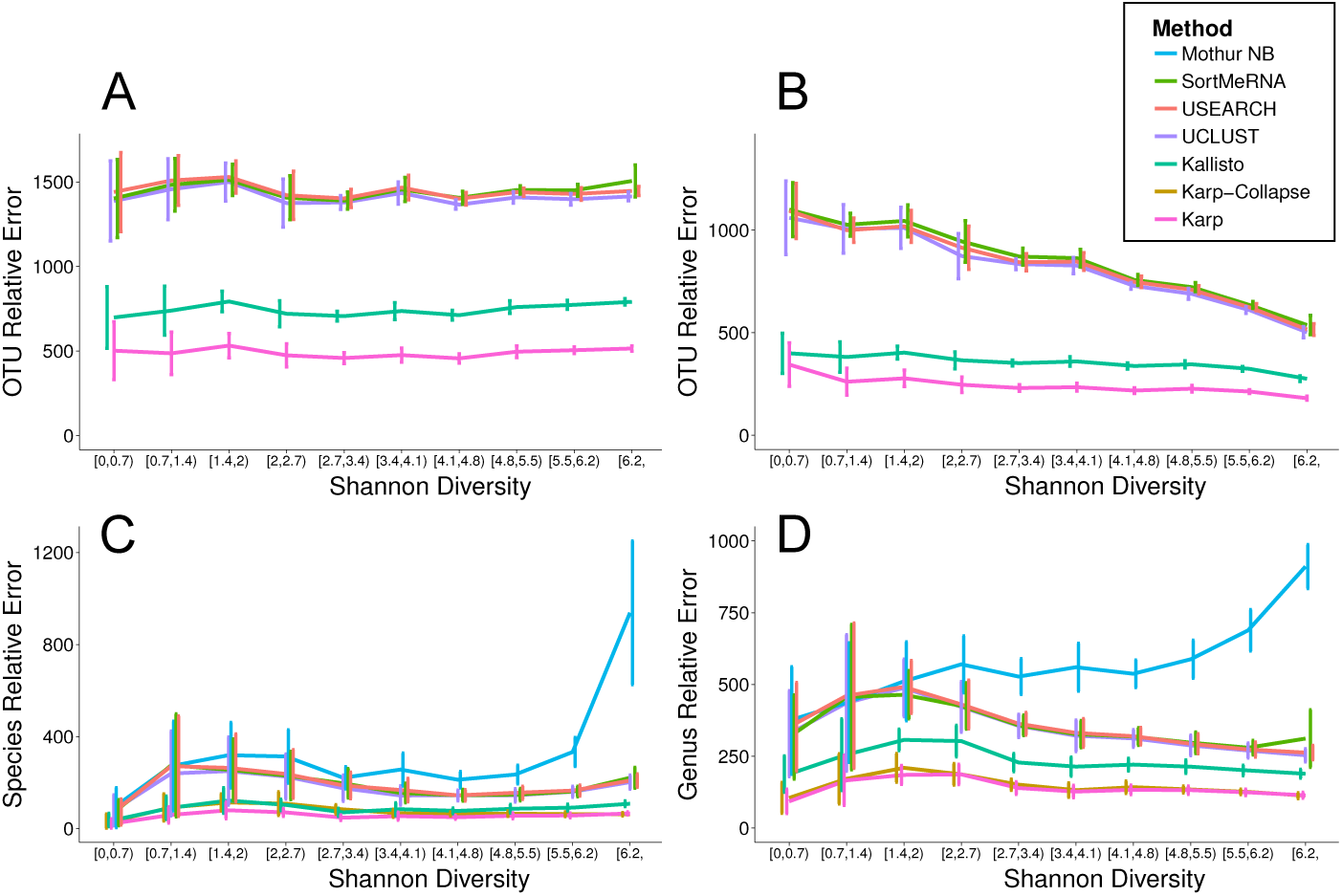
The average absolute error with 95% confidence intervals from simulated samples of 1 × 10^6^ 75bp reads, with every simulated dataset having a unique mix of 1,000 reference haplotypes drawn from the GreenGenes database. Each colored line represents a different classification method, including Karp, Kallisto, UCLUST, USEARCH, SortMeRNA, and the Naive Bayes method implemented in Mothur. Error refers to the average relative error (AVGRE): the difference between the true number of reads for each reference haplotype present in the simulated data and the number classified by each method, if each method had classified every read in the data. (A) Total OTU-level error for taxa with > 1 read present. (B) OTU-level error for taxa with frequency > 0.1%. (C) Species-level error. (D) Genus-level error.

Many reference haplotypes share the same taxonomic label, and it is possible researchers would be interested in hypothesis at the level of genus or species rather than individual references. We aggregated counts for references with identical labels and again compared with the truth in our simulated samples. When we compared estimates at the level of both species and genus, on average the full Karp algorithm was the most accurate method for samples with a broad range of diversity (Shannon diversity < 6.2) while Karp-collapse performed best in the most diverse samples (Shannon diversity > 6.2) (Figures 2C and 2D). At higher level taxonomic classifications Karp remained the most accurate method (Supplementary Figures S7 and S8).

The difference in classification error observed here is relevant for downstream analysis. When we calculated summary statistics using OTUs with frequencies > 0.1% in 100 independent simulated samples, Karp’s estimates were on average closer to the truth than either Kallisto or UCLUST (Table 1). For Simpson Diversity, Karp’s estimate was within 10% of the actual value for 44% of samples, compared with 32% of samples for Kallisto, and only 2% of samples with UCLUST. Karp’s estimate of Simpson Diversity fell within 25% of the actual value in 82% of samples, with Kallisto this figure was 62%, and UCLUST was 18%.

**Table 1:** Summaries of microbiome data were calculated from 100 simulated samples containing different mixtures of 1,000 references. Only reference haplotypes with frequencies > 0.1% were used to calculate the statistics. In each sample the absolute value of the difference between the actual statistic and that estimated by Karp, Kallisto, and UCLUST was calculated. The group-wise Beta Diversity value was a single estimate from all 100 samples; it is not an average and therefore there is no standard error.

|  | Individual | Pairwise |  | Group |
| --- | --- | --- | --- | --- |
| Statistic | Simpson Diversity | $^2D$ Beta Diversity | Bray-Curtis Dissimilarity | $^2D$ Beta Diversity |
| Actual Values | 0.002 - 0.9 | 0.44 - 0.87 | 0.40 - 1.0 | 0.025 |
|  | Average Absolute Difference (Standard Error) |  |  |  |
| Karp | 0.014 (0.029) | 0.029 (0.035) | 0.009 (0.018) | 0.002 |
| Kallisto | 0.015 (0.025) | 0.037 (0.046) | 0.012 (0.023) | 0.002 |
| UCLUST | 0.079 (0.13) | 0.093 (0.090) | 0.026 (0.043) | 0.008 |

In addition to 75bp single-end reads, we simulated and classified samples with longer paired-end reads. We simulated and classified 130 samples with 151bp paired-end reads and 170 samples with 301bp paired-end reads. On average, when we compared estimates for references with frequency > 0.1% in the 151bp paired-end samples Kallisto was the most accurate method for datasets with very low Shannon Diversity (< 0.7), Karp and Kallisto performed nearly identically for samples with low to moderate Shannon Diversity (0.7 – 3.4), and Karp had the lowest error when Shannon Diversity was high (> 3.4) (Figure 3A). When we aggregated counts for OTUs with identical taxonomic labels and compared the abundance estimates of species with frequency > 0.1%, Karp had the lowest average errors (50%, 84%, and 94% less than Kallisto, UCLUST, and the Wang *et al*. Naive Bayes respectively, Figure 3B). Of the three read lengths examined, Karp’s advantage was greatest for the 301bp paired-end reads. For these reads Kallisto’s strict pseudoalignment threshold struggled to make assignments, and on average classified only 3.8% of reads (versus 53% with Karp). Note that Kallisto’s performance could be improved by subsampling shorter regions from the longer reads, although this would be removing information that Karp is currently using to assign reads. Also, the Naive Bayes classification could not be computed for the 301bp paired-end reads with the computational resources available for this project and was therefore not compared. In references with frequency > 0.1% Karp was on average the most accurate method across the entire range of Shannon Diversity (errors 90% smaller than Kallisto, 62% smaller than UCLUST, Figure 3C).

**Figure 3:**
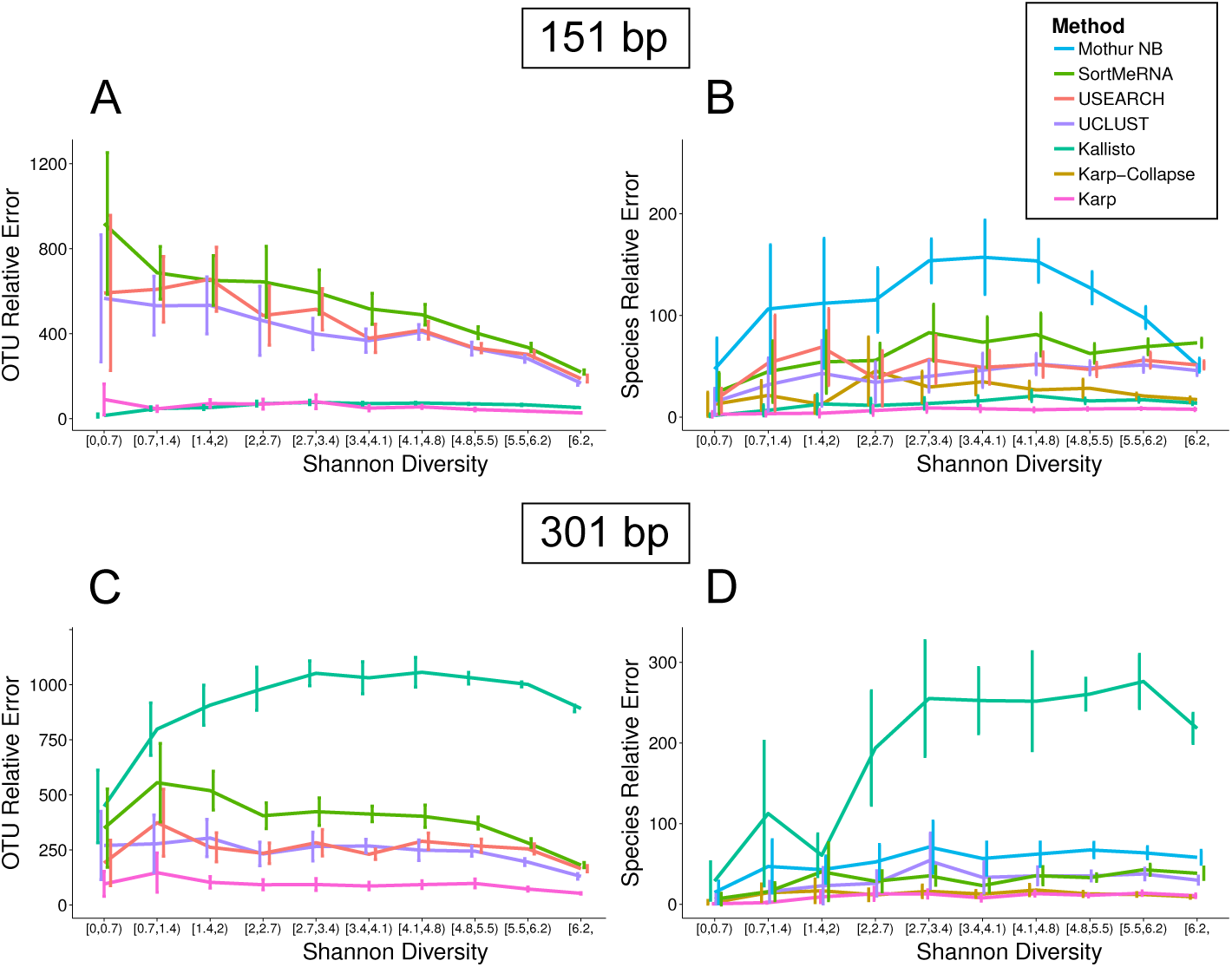
Average relative errror (AVGRE) with 95% confidence intervals from the taxonomic classification of simulated samples with 151bp paired-end and 301bp paired-end reads. Taxonomy was classified using Karp, Kallisto, UCLUST, USEARCH, SortMeRNA, and the Naive Bayes method implemented in Mothur. (A) OTU-level error in 130 samples of 151bp paired-end reads for OTUs with frequencies > 0.1% (B) Species-level error in 130 samples of 151bp paired-end reads for species with frequencies > 0.1%. (C) OTU-level error in 170 301bp paired-end samples for OTUs with frequencies > 0.1%. The strict pseudoalignment threshold for Kallisto reduced the number of reads classified and increased the errors in its estimates. (D) Species-level error in 170 301bp paired-end samples for species with frequencies > 0.1%. For computational reasons we were unable to calculate Naive Bayes estimates for the 301bp samples.

In microbiome classification problems it is not uncommon to have taxa present in sequenced samples that are absent from reference databases. We tested the robustness of Karp under this scenario with simulated samples containing reads from haplotypes removed from the reference databases used for classification. For each of one phylum (Acidobacteria), one order (Pseudomonadales), and one genus (Clostridiisalibacter) we simulated 10 independent datasets where 50% of reads originated from 3 different members of each taxon and created copies of the GreenGenes database were the reference sequences for every member was removed. We classified the simulated data with both the reduced databases and the full GreenGenes to measure how much the absence of relevant references impacted estimate accuracy. Karp, Kallisto, and UCLUST were all less accurate when classifying samples using the reduced databases rather than the full database (Figure 4). Under all scenarios Karp remained the most accurate method, and in the case of the phylum Acidobacteria and genus Clostridiisalibacter Karp’s classification using the reduced reference database was more accurate than UCLUST using the full reference database.

**Figure 4:**
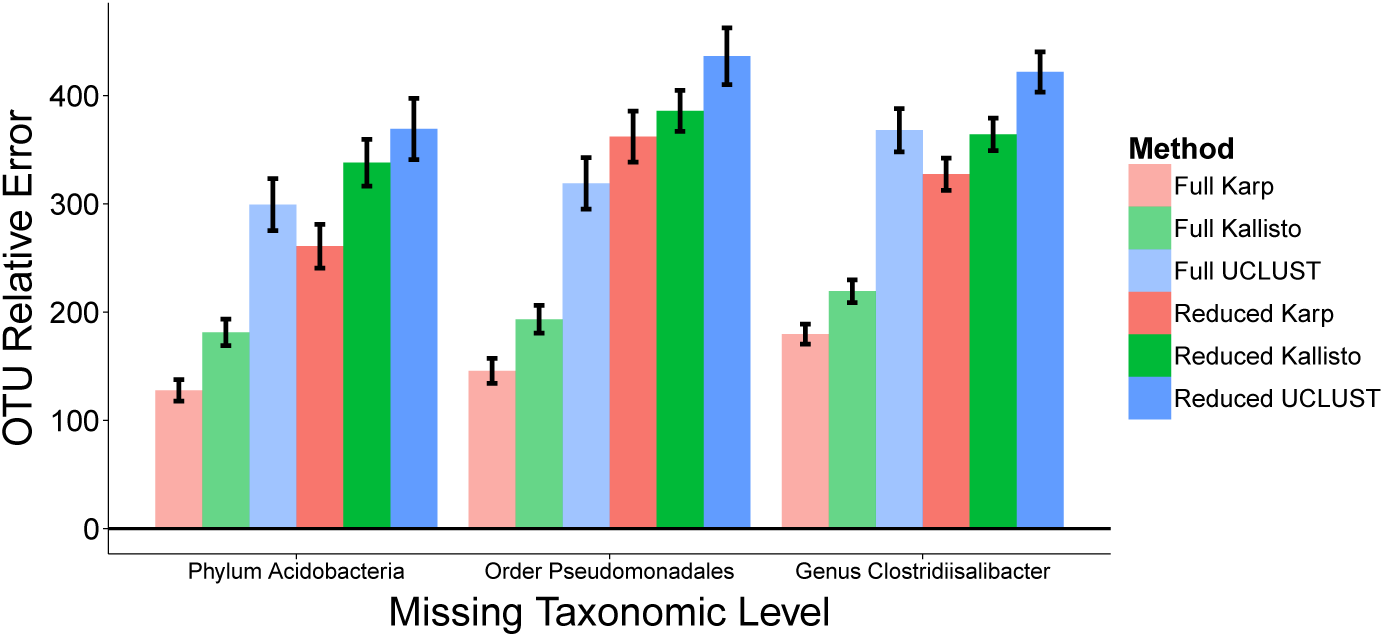
Accuracy when the reference database used for classification is missing taxa found in the sample. For each of one phylum (Acidobacteria), one order (Pseudomonadales), and one genus (Clostridiisalibacter), 10 samples were simulated where 50% of the reads originated from the noted taxa. Each sample was classified with the full GreenGenes database and also a reduced version of the database lacking all members of the taxa which had been used to simulate the sample. The accuracy of estimates by Karp, Kallisto, and UCLUST for the 50% of the samples that did not originate from the absent taxa were compared with their true frequencies. Black bars give 95% confidence intervals.

The model that underpins Karp relies on knowing the probability of a sequencing error at a given position in a read. Our work assumes that the base-quality scores are accurate estimates for the probability of sequencing error. In real data it has been recognized that base-quality scores are not always accurate, leading to the development of methods to empirically recalibrate base-quality m using known monomorphic sites. With pooled microbiome samples this recal-ibration is complicated (possibly requiring spike-ins of known sequence during the experiment or alignment to conserved reference sequence), so we explored how differences between the expected sequencing error rate as represented by the base-quality scores and the actual sequencing error rate effect Karp’s accuracy. Under a model where the actual probability of a sequencing error was 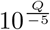 for *Q* quality score Q, rather than the 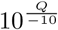 assumed by our model, Karp was still on average more accurate than Kallisto or UCLUST/USEARCH (errors 12.9% smaller than Kallisto and 64.2% smaller than UCLUST, Supplementary Figure SS6A). When errors actually occurred uniformly at 1% of bases, grossly violating Karp’s model, it was still the most accurate method (errors 11.5% smaller than Kallisto, 62.8% smaller than UCLUST, Supplementary Figure SS6B).

The increased accuracy of Karp comes at some computational cost, especially relative to Kallisto, however it is still quite feasible for modern data. Table 2 compares the performance of the methods while classifying samples with either 10^6^ 75bp single-end, 151bp paired-end, or 301bp paired-end reads using 12 cores, and in all cases even the full mode of Karp requires < 3 hours. Karp was run with default settings, in both full and collapse mode. For Karp and Kallisto the 75bp reads require longer to classify than the 151bp reads due to the larger number of multiply-mapped reads with the shorter length.

**Table 2:** Computational requirements and speed of Karp, Kallisto, UCLUST, USEARCH61, SortMeRNA, and the Wang *et al*. (2007) Naive Bayes using Mothur. All programs were run using 12 multi-threaded cores except Mothur. Mothur’s memory requirements scale with the number of cores used, and in order to keep memory <16GB we limited it to 4 cores. The values for UCLUST and USEARCH give the time to assign taxonomy, generally with these methods reads are clustered before taxonomy is assigned and the value in parenthesis gives the time to first cluster and then assign taxonomy.

| Method | Time (Minutes) |  |  | Max Memory |
| --- | --- | --- | --- | --- |
|  | 75bp | 151bp | 301bp |  |
| Karp Full | 161.9 | 24.7 | 80.4 | 10 GB |
| Karp Collapse | 36.0 | 16.1 | 74.5 | 10 GB |
| Kallisto | 4.5 | 2.6 | 1.3 | 10 GB |
| UCLUST | 87.4 (146.6) | 18.9 (67.1) | 3.9 (55.9) | 4 GB |
| USEARCH61 | 9.6 (27.7) | 1.4 (10.0) | 0.7 (7.1) | 4 GB |
| SortMeRNA | 37.3 | 22.3 | 29.6 | 4 GB |
| Mothur Naive Bayes* | 502.4 | 1578.2 | NA | 16 GB |

\*limited to 4 cores

### 3.2 Performance assessment in real 16S rRNA data

We classified 368 16S rRNA samples collected from the shoes, phones, and floors of two study participants using Karp, Kallisto, and UCLUST. For each classification method we subsampled without replacement 25,000 reads from each sample, either from individual references or else after aggregating counts within taxonomic labels, and then performed several analyses. For robustness, we performed the subsampling 10 times for each method and analysis. First, we used the random forest classification method with 1,000 trees to classify subsets of the data. We classified the shoe samples as coming from person 1 or person 2. We did the same with the phone samples, and then within each individual we classified the phone samples as either from the front or back of their phone, and their shoe samples as coming from the front left, front right, back left, or back right. In these analyses we measured the classification error using the known identity of each sample (Table 3). When we aggregated counts by taxonomic label there were not enough reads at the species level to subsample, so we performed the classification with genus-level labels. Error rates were lower when classification was done using individual references rather than counts aggregated by genera. The error rates for classifying the shoe surfaces from person 2’s samples were greater than the baseline error rate (if every sample had been assigned the most common label), suggesting there was not power to perform this classification.

**Table 3:**
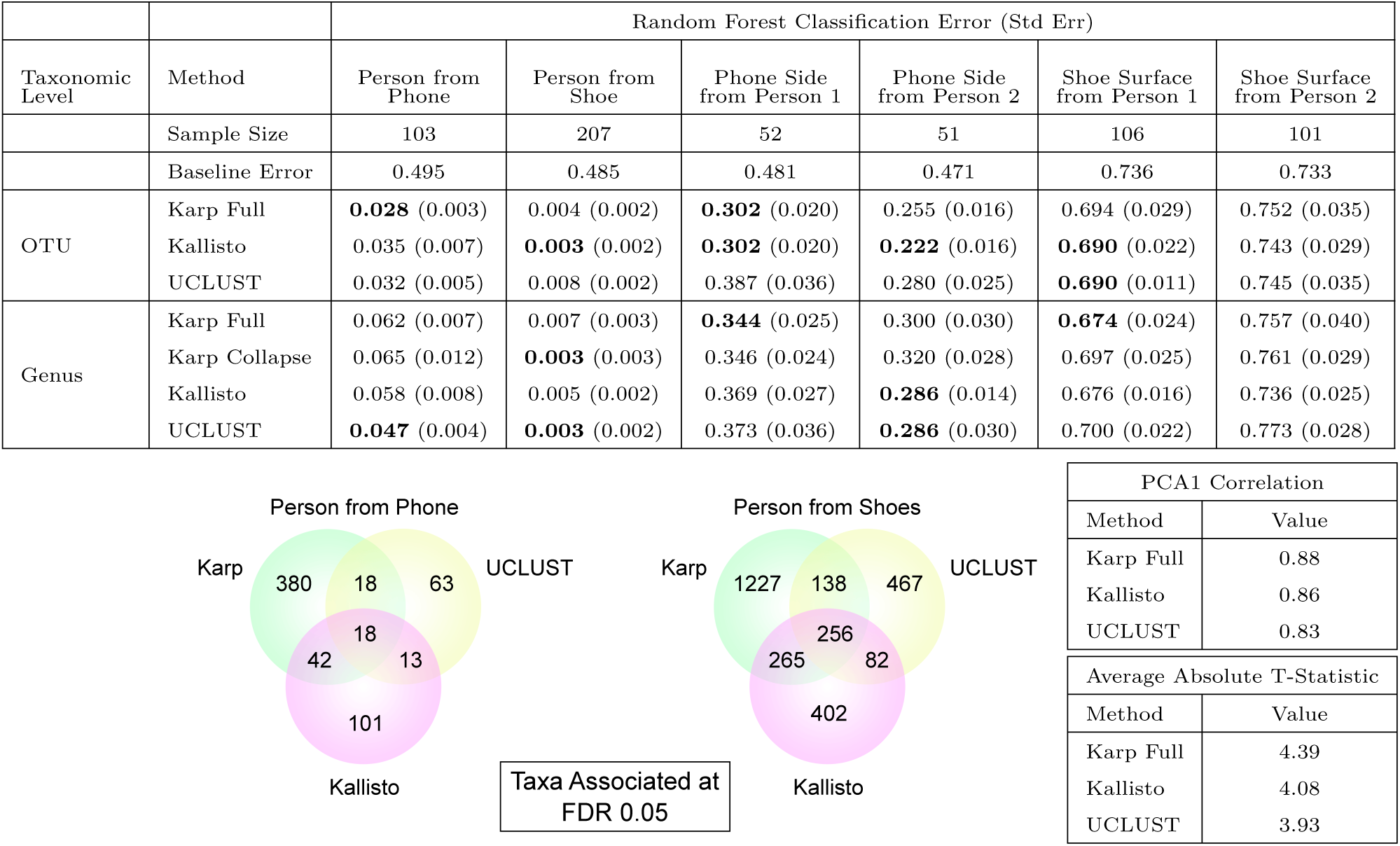
After samples were classified with Karp, UCLUST, and Kallisto 25,000 reads were subsampled 10 times and three analyses were performed. (Top) We used random forests with 1,000 trees to classify the origin of the samples. The average classification error for each method was recorded from 10 independent subsamplings. Bold text indicates the best performing method within a category. (Bottom) Separately within the phone and shoe samples we tested for differences in the mean abundance of taxa between individuals 1 and 2 using Welch’s t-tests. We tested taxa with at least 250 reads observed across all samples. The Venn diagrams display the number of taxa observed to be different at an FDR of 0.05 in all 10 subsampled matrices for each method. The table on the bottom right also includes the average absolute value of the t-statistics with an FDR < 0.05 from each method. (Bottom Right) We performed a principal components decomposition on the subsampled matrices and looked for correlation in PCA1 between shoe and floor samples collected side by side.

Next, we performed a PCA decomposition on matrices with 25,000 reads subsampled from each of the floor and shoe samples. From this we calculated the average correlation between PCA1 for each floor sample and the shoe samples collected at the same location and time (Table 3). Karp had the greatest average correlation (0.88), next was Kallisto (0.86), and finally UCLUST (0.83).

Finally, we tested for differences in the mean abundance of taxa between person 1 and person 2 using Welchs’ t-tests. We tested OTUs with at least 250 observed reads across all samples, and tested the phone and shoe samples separately. When we looked at taxa that varied significantly between individuals with a false discovery rate (FDR) < 0.05 in all 10 matrices of subsampled reads, Karp detected the most differentiated taxa (458 phone, 1,886 shoes) compared with UCLUST (112 phone, 943 shoes), and Kallisto (174 phone, 1,005 shoes) (Table 3). The average strength of association as measured by the absolute size of the t-statistic was also larger in the Karp analysis (4.39), than for Kallisto (4.08) or UCLUST (3.93).

## 4. Discussion

In both simulations and real 16S data we have shown that Karp is an accurate and computationally feasible method for estimating the relative frequencies of contributing members in a pooled DNA sample. Although not as fast as some alternatives, Karp’s superior accuracy across the tested range of read lengths, taxonomic levels, and absent references makes a strong case for its adoption.

Although our work here has focused on applying Karp in the context of 16S microbiome experiments, its potential uses extend to most closed-reference classification problems. Pooled DNA experiments are common in many fields. The identification and estimation of contributor abundance in whole genome metagenomics, pooled sequencing of data from artificial selection experiments, and RNA isoform identification are all possible with Karp.

In order to perform well Karp needs sequencing reads that contain enough information to distinguish between the reference sequences. Our simulations, where Karp outperformed the alternative methods convincingly, used sequence from the entire 16S rRNA gene, which had sufficient information to distinguish between almost all closely related references. In the Lax *et al*. (2015) data, and frequently in 16S sequencing projects, a limited number of the gene’s hyper-variable regions are sequenced. In such a limited reference region, many closely related haplotypes are identical or nearly so, and there is little information to distinguish between them. Here the difference between methods that probabilistically assign reads to references, like Karp and Kallisto, and those that make a hard assignment, like UCLUST or USEARCH, can arise. With Karp, when references are nearly identical they will receive nearly equal probability weights from each read that maps to them, and the result will be many closely related references at low frequencies. With UCLUST or other similarity score methods, the references are sorted and the first of the closely related references to appear in the sorting order will be assigned all or nearly all the references, regardless of if it is the actual contributing organism. The truth in this case, is that the sequencing data does not contain enough information to accurately distinguish between the references, and both methods end up at sub-optimal, albeit different solutions. Under such conditions researches need to have a realistic expectation of what they can resolve in their data, and it is likely that inferences of higher-level taxonomic abundances rather than individual references are more likely to be robust.

Current experimental protocols and downstream clustering algorithms make using a single hypervariable region in the 16S gene a standard approach. A single hypervariable region is short enough that it is rare for reads from a single organism to form multiple OTUs during clustering. However, it is important to understand that the sequencing of a smaller reference costs researchers information that could make it possible to improve quantification accuracy and distinguish between closely related references. Our simulations suggest experimenters could benefit substantially from sequencing more of the 16S gene than is often presently used.

In addition to k-mer length, Karp users can adjust the thresholds for minimum frequency during the update step of the EM algorithm and the likelihood filter z-score. While results are often relatively invariant across a broad range of threshold values (Supplementary Figures S1, S2, S3, S4), avoiding extreme threshold values can improve classification accuracy substantially. Practical guidance for setting the thresholds is given in supplementary section 7.3. It is worth noting that Karp’s tuning parameters influence performance as well as accuracy, so choosing optimum values can improve not just accuracy but experimental run time as well.

In both ecology and human health a greater understanding of the microbiome promises medical and scientific breakthroughs. Modern sequencing technology gives us unprecedented access to these microbial communities, but only if we can correctly interpret the pooled DNA that sequencing generates can we hope to make significant progress. Towards that end, Karp provides a novel combination of speed and accuracy that makes it uniquely suited for scientists seeking to make the most out of their samples.

## 5 Acknowledgments

This work was supported by NIH/NHGRI R01 HG007089. We would like to thank Lior Pachter for helpful suggestions about pseudoaligning and improving program performance. Additonal thanks are due to Katie Iguarta and the Ober Lab as well as Simon Lax and the Gilbert Lab at University of Chicago for sharing technical expertise and data for this study. Chaoxing Dai wrote a C++ version of the R-package SQUAREM that provided a template for our own implementation in Karp. Emily Davenport provided helpful feedback on the manuscript. This work was completed in part with resources provided by the University of Chicago Research Computing Center.

## 7 Supplement

### 7.1 EM algorithm

For a pooled sample of reads *r* with *r* ∈ 1,…,*N*, if we observed which reference haplotypes the reads in our sample originated from, *η*, and we assumed that conditional on the frequencies 𝓕 the query reads are independent, it would be possible to calculate the maximum likelihood estimate 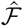 using the complete data likelihood, which has the form

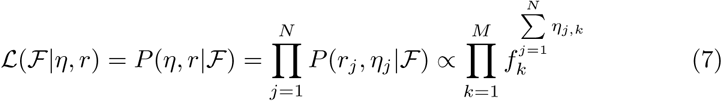

In actuality, we observe the reads but the reference haplotypes that they originate from are unobserved. To estimate 𝓕 we therefore employ an EM algorithm. Briefly, the E-step of our procedure can be written

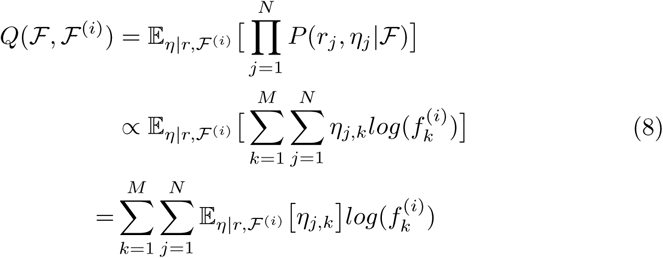

where

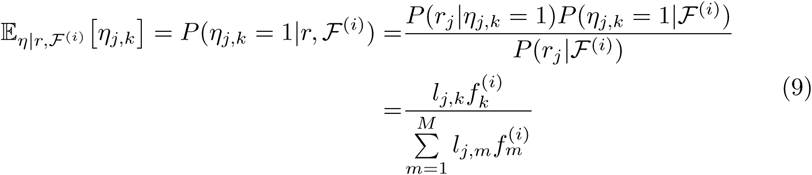

The M-step directly follows from the form of our likelihood, and the algorithm updates the estimates of 𝓕 until convergence according to

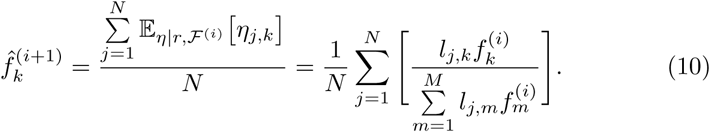

### 7.2 Likelihood filter

When we classify pooled microbiome data it is likely that some reads originate from taxa that are absent from our reference database. Filtering these reads improves the accuracy of frequency estimates for the taxa that are present. Karp uses a likelihood based filter that was first published and validated in Kessner *et al*. (2013).

Given a set of query reads with their corresponding base-quality scores, we can calculate the mean and variance for the distribution of likelihood values that would result if every query read were aligned to the actual reference that gave rise to it, such that every mismatch was the result of sequencing error. This calculation requires only the query read base-quality scores, not the actual reads or a reference database, and is carried out before Karp begins pseudoalignment.

Recalling the notation of section 2.3, a read of length *L*, has bases *r*_[0]_, *r*_[1]_, …,*r*_[*L*]_ and corresponding base-quality scores *q*_[0]_,*q*_[1]_, …,*q*_[*L*]_. If each read *r* originated from a reference *h*, our goal is to calculate 𝔼 [*log*(*P*(*r*|*q*, *h*))] and *Var* [*log*(*P*(*r*|*q*, *h*))].

First, for each position *i* ∈ 1, …, *L* define the empirical distribution of base-quality scores, *Q*_[*i*]_, in a sample of *N* reads by

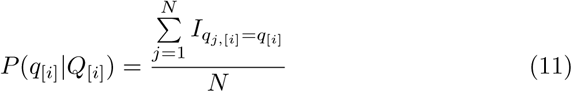

where *I* is an indicator function and *q*_*j*,[*i*]_ is the base-quality score at position *i* on read *j*. This distribution is independent of *h*.

Assuming that each position along a read is independent we can write:

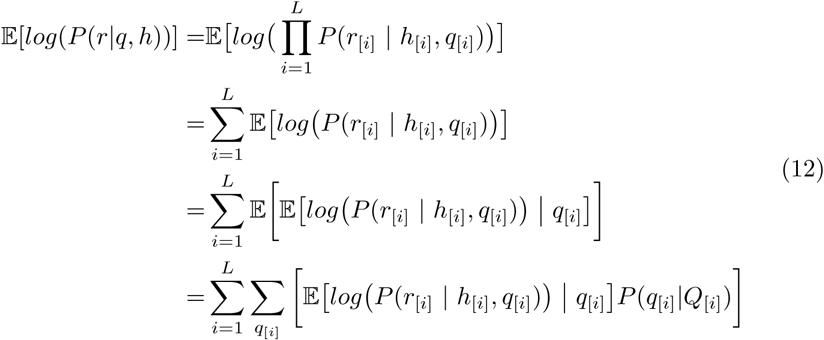

For each position *i* the probability of sequencing error is a known function of the base-quality score, *ɛ*(*q*_{*i*]_). Karp assumes Phred scaled base-quality scores (with options for Phred+33 or Phred+64), where *ɛ*(*q*_{*i*]_) = 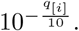 Using *ɛ*(*q*_{*i*]_) and equation 2 we can write the conditional expectation as:

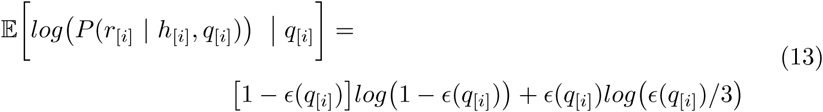

Note that this expression does not depend on *h*_[*i*]_ or *r*_[*j*]_. By combining equations 11, 12, and 13 we have an expression for 𝔼[*log*(*P*(*r*|*q*, *h*))]. To calculate 814 *Var* [*log*(*P*(*r*|*q*, *h*))] we again use the assumption that bases are independent and write:

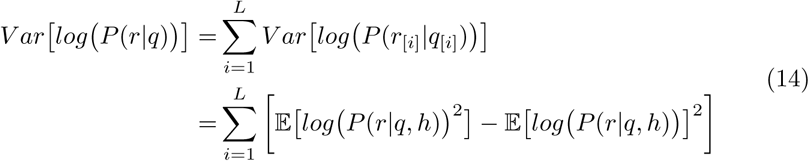

The likelihood filter is applied after the query reads have been locally aligned to the reference database and the corresponding likelihood values have been 818 determined. Then, a z-score is computed for each query read using its largest likelihood value and the mean and variance of the “null” likelihood distribution (Equations 13 and 14). If this z-score is too low it is evidence that the true reference that the read originated from is absent from the database, and correspondingly the read is removed.

### 7.3 Effect of Karp tuning parameters on run-time and accuracy

While accuracy is largely similar across a range of values, understanding when adjusting the minimum EM update frequency or the z-score could improve results is important for Karp users. Setting the minimum EM frequency threshold too high causes the removal of real references present in the sample, while setting it too low can cause spurious references to be included in the final solution. In situations with enough information to distinguish between closely related references, for example if the entire 16S gene sequence has been sequenced, a greater frequency threshold can yield more accurate solutions (Figure S3). Under such conditions threshold values on the order of Karp’s default (1/Number of reads) are often appropriate. Alternately, where only limited information exists, for example if a single hypervariable region has been sequenced, lower thresholds can give more optimal solutions (Figure S4 and Tables S1 and S2). This is because a lower threshold avoids removing organisms truly in the sample that have had their reads spread across closely related taxa, each with a fraction of the true organism’s frequency. With limited information setting the minimum frequency an order of magnitude lower than the Karp default (i.e. 840 1/(10 _*_ Number of reads)) can yield better results.

**Figure S1:**
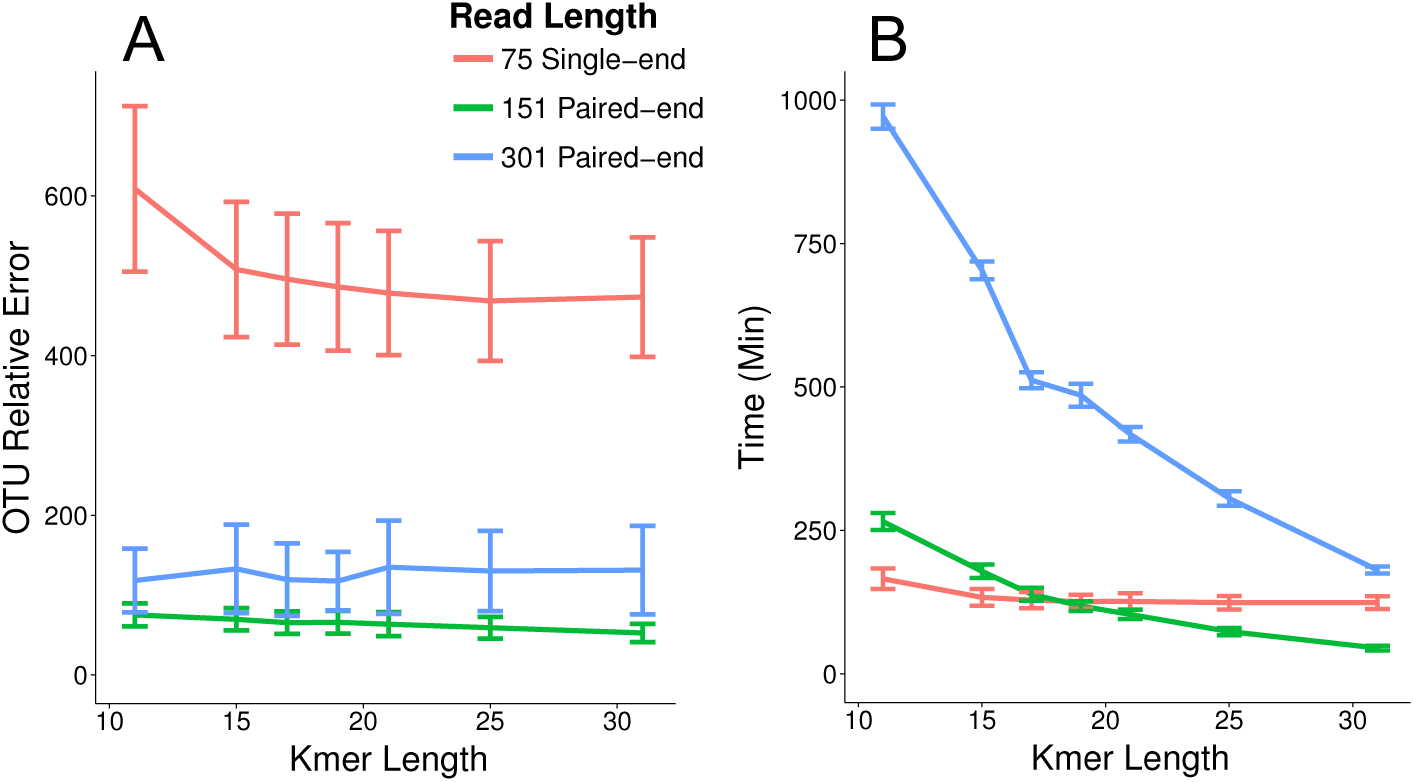
Impact of k-mer length on Karp performance. Pseudoalignment indexes constructed using different k-mer lengths were used to classify 30 previously analyzed samples selected to cover a full range of Shannon Diversities. For each of 75bp, 151bp, and 301bp reads 10 samples were analyzed. (A) The average error values with 95% confidence intervals for each read length. (B) Average run times using 12-cores in parallel.

For the z-score likelihood filter, Karp estimates the mean and standard de-viation using the distribution of base-quality scores present in the data being classified. Thus, the quality of the data plays a role in determining the best threshold to use. Karp includes an option to output the distribution of maximum likelihood scores for a sample. Outputting these scores for a few samples, and comparing them with the z-score values output in Karp’s log files is a good way to determine if the default threshold is too strict or lenient for a particular experiment. If the threshold is falling too near the median value of the real likelihoods, lowering it may improve accuracy by retaining more reads. If the threshold falls far outside the actual distribution of read likelihoods, setting it to a greater value could improve its ability to filter our reads from references absent from the reference database.

**Figure S2:**
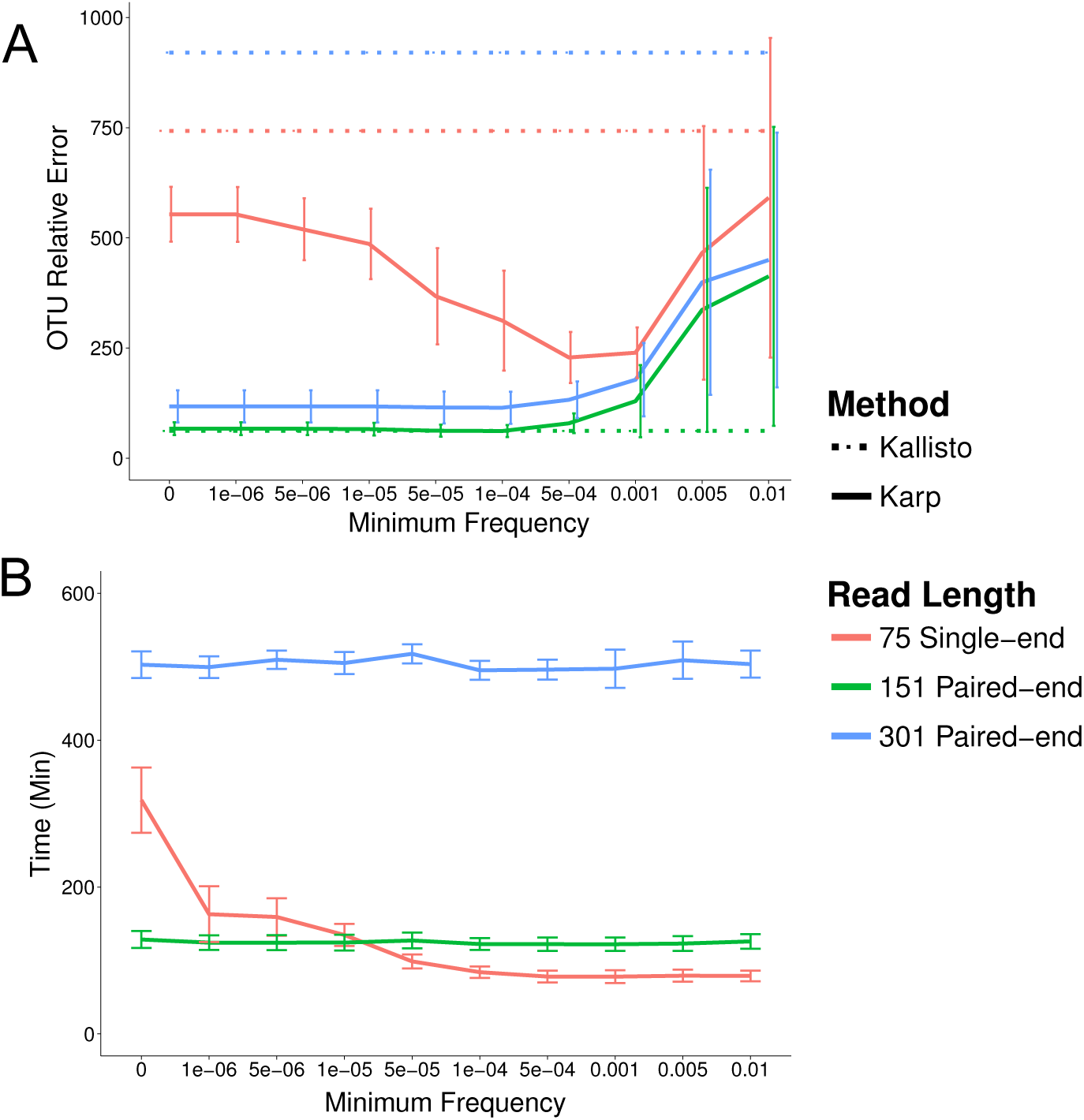
Karp uses an EM algorithm to estimate the relative frequencies of reference haplotypes in a pooled sample. During the EM process a minimum frequency threshold can be applied that removes references with frequencies below this threshold. Set at a low frequency, the threshold helps remove spurious findings and improves accuracy, particularly for shorter reads. At higher frequencies the threshold removes references actually present in the sample and lowers accuracy. In this figure different thresholds are applied during classification of 30 previously analyzed samples selected to cover a full range of Shannon Diversities. K-mers of length 19 were used for these analyses. For lengths of 75bp, 151bp, and 301bp 10 samples were analyzed. (A) The average error values with 95% confidence intervals for each read length. (B) Average run times using 12-cores in parallel. For shorter reads, increasing the threshold reduces the number of EM iterations required to converge and decreases run-time.

**Figure S3:**
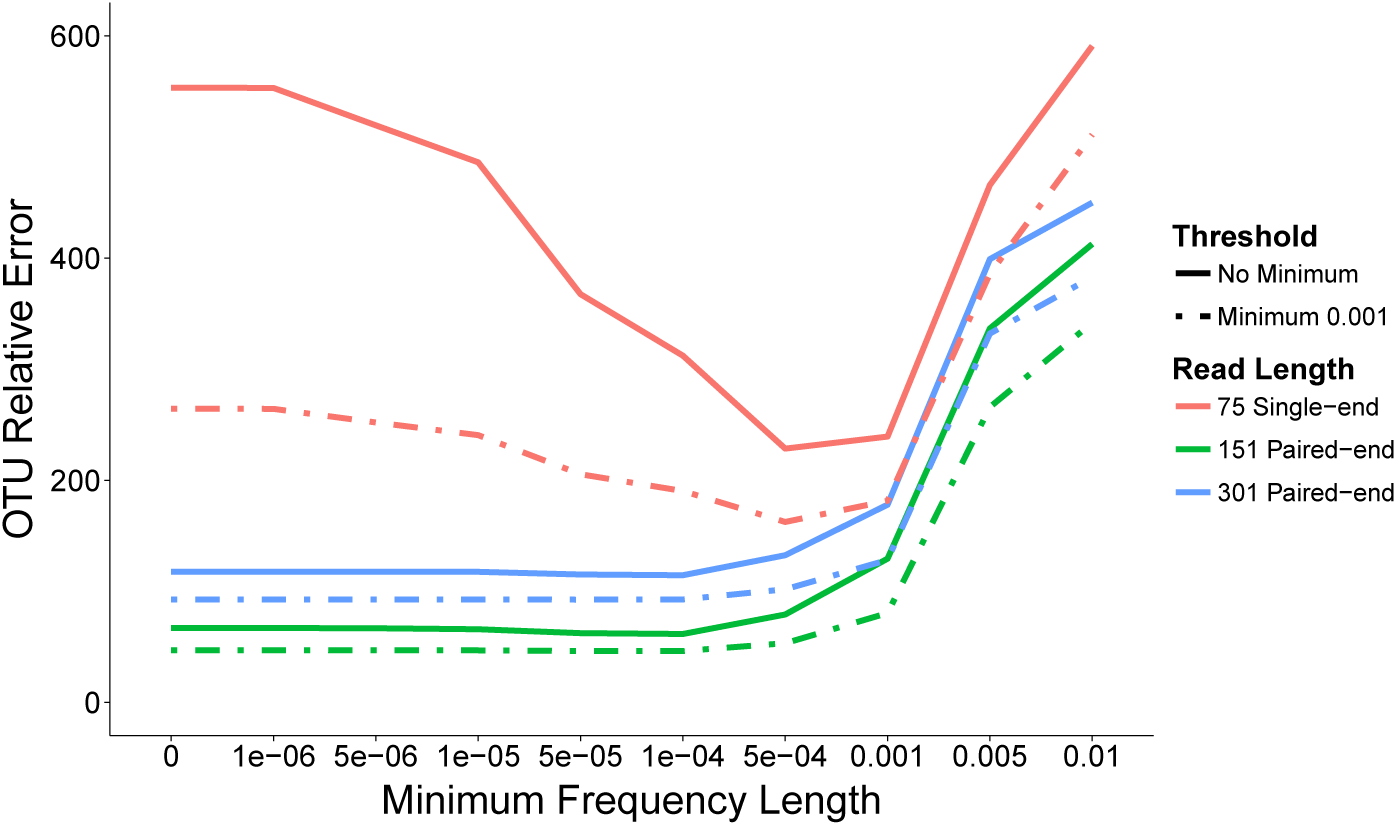
The impact of the EM frequency threshold is smaller when analyzing error in the estimates of more common references. Solid lines present the error calculated using all references classified, dashed lines give the error when only references with an actual or estimated frequency above > 0.1%, a cut-off used frequently in this study. In such cases the chosen frequency threshold is less important.

**Table S1:** In the Lax *et al*. (2015) shoe and phone data, we evaluated the impact of a range of Karp tuning parameter values (EM algorithm minimum frequency and maximum likelihood z-score) on random forest classification error. This table reports how often a given EM minimum frequency had the lowest (or tied for the lowest) error rate for a particular analysis in figure S4. Higher counts reflect better performance, the maximum count in row 1 would be 12, and in row 2 would be 8. Particularly for the full version of Karp, setting a lower minimum frequency resulted in lower classification error rates.

|  | Parameters with Lowest Classification Error |  |  |  |  |  |  |  |
| --- | --- | --- | --- | --- | --- | --- | --- | --- |
| Minimum frequency | $1 \times 10^{-2}$ | $1 \times 10^{-3}$ | $1 \times 10^{-4}$ | $1 \times 10^{-5}$ | $1 \times 10^{-6}$ | $5 \times 10^{-7}$ | $1 \times 10^{-7}$ | $1 \times 10^{-8}$ |
| Karp Full | 0 | 0 | 1 | 2 | 1 | 0 | 3 | 5 |
| Karp Collapse | 1 | 1 | 0 | 0 | 1 | 1 | 2 | 2 |

**Table S2:** In Lax et al. (2015) shoe and phone data, we evaluated the impact of a range of Karp tuning parameter values (EM algorithm minimum frequency and maximum likelihood z-score) on classification error from randomForest classification. This table reports how often a given maximum likelihood z-score cutoff had the lowest (or tied for the lowest) error rate for a particular analysis in figure S4. In both modes of Karp more lenient thresholds resulted in lower classification error rates.

|  | Parameters with Lowest Classification Error |  |  |  |  |  |  |  |  |
| --- | --- | --- | --- | --- | --- | --- | --- | --- | --- |
| Likelihood z-score | -0.5 | -1 | -1.5 | -2 | -3 | -4 | -5 | -6 | -7 |
| Karp Full | 1 | 1 | 2 | 2 | 0 | 1 | 0 | 4 | 3 |
| Karp Collapse | 1 | 1 | 1 | 0 | 0 | 0 | 1 | 1 | 4 |

**Figure S4:**
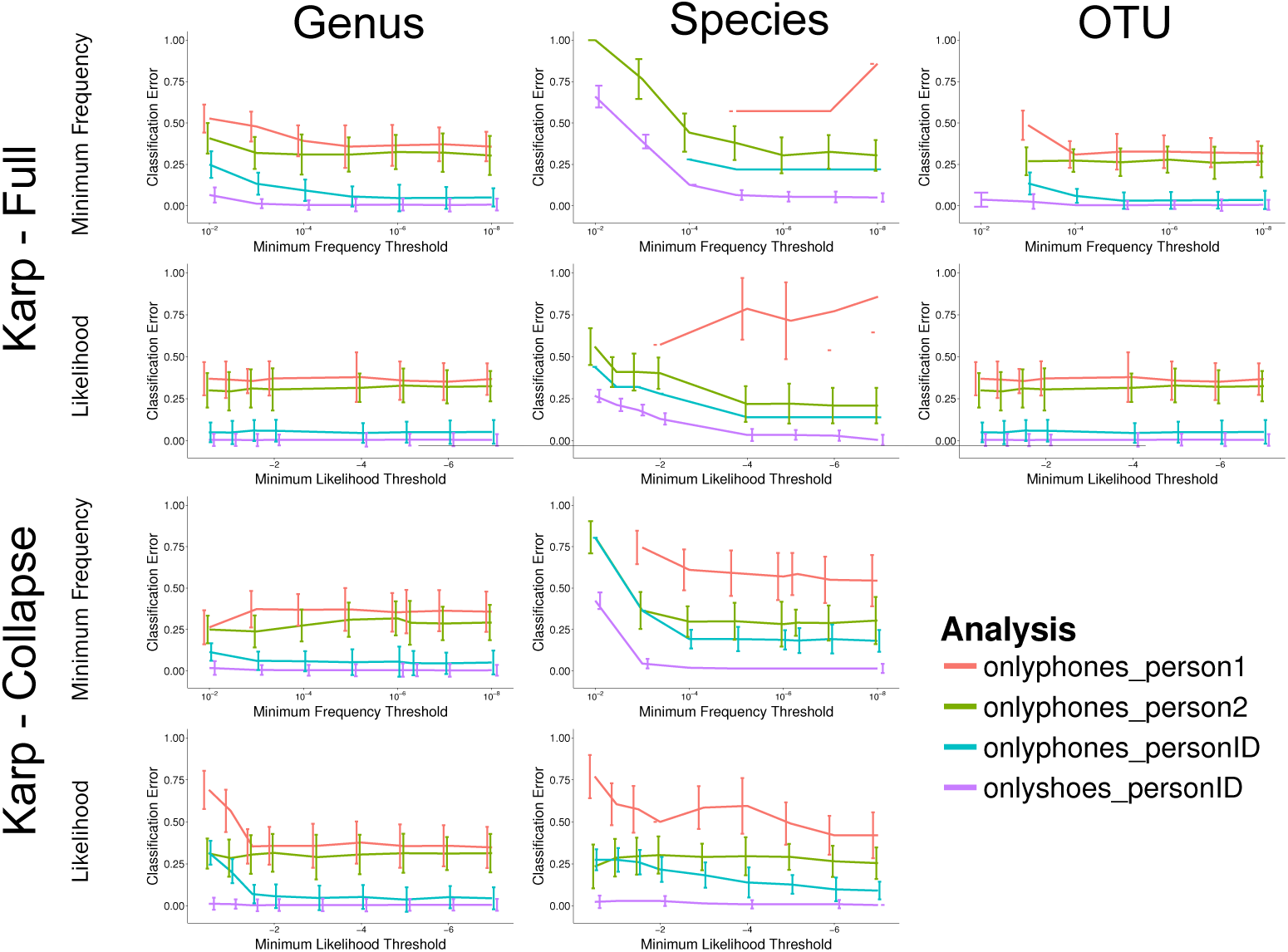
Evaluating impact of Karp tuning parameters (EM frequency threshold and maximum likelihood z-score) on shoe and phone data from Lax et al.(2015). We classified each sample using a range of tuning parameters and then performed random forest classification with 10,000 subsampled reads per sample and 1000 trees. We used both the Full and Collapse mode of Karp. Each colored line represents a different analysis, and the bars give 95% confidence intervals based on 10 replicates. When a parameter setting failed to assign taxonomy to enough reads to classify at a subsampled depth of 10,000 reads where other parameter settings successfully quantified to that depth, it was counted as an error for the purpose of random forest classification.

**Figure S5:**
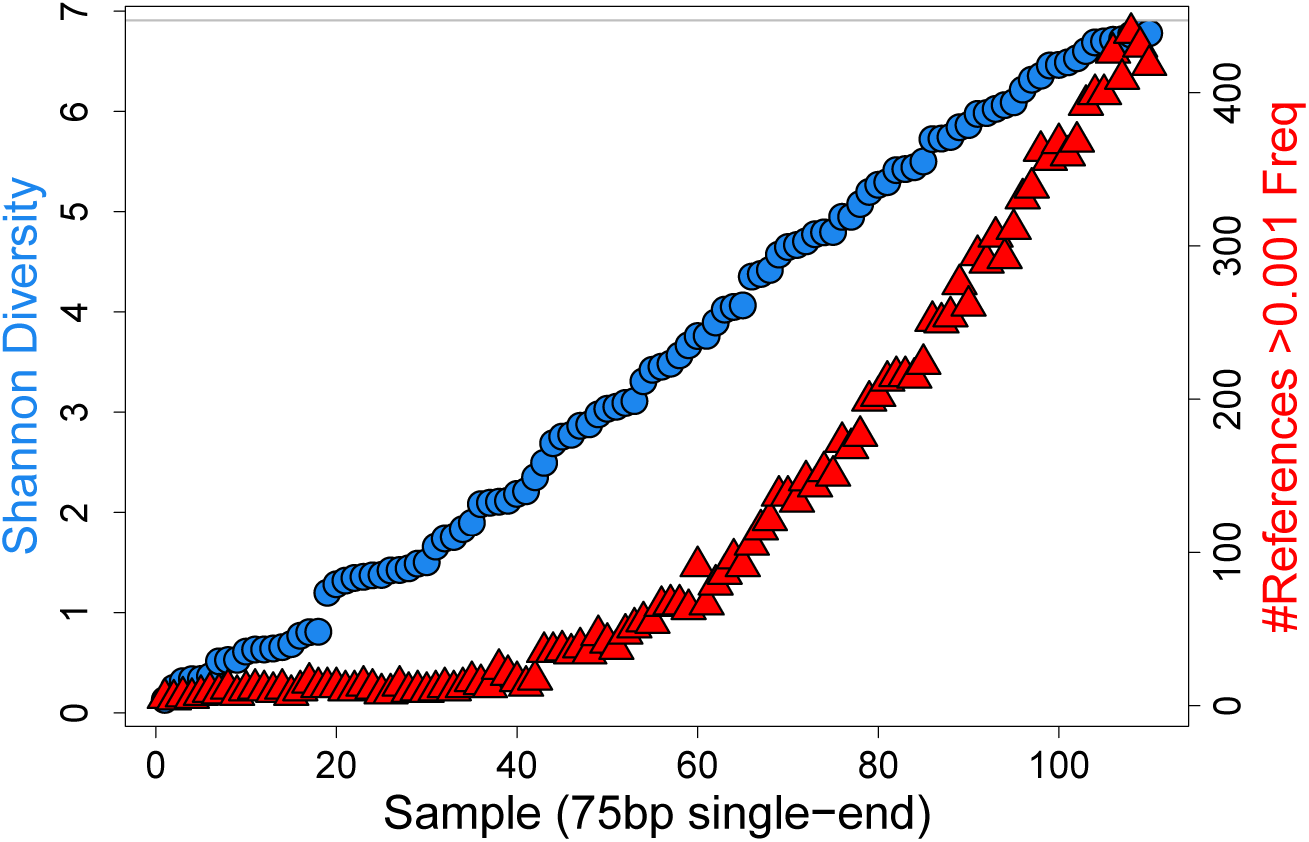
Each simulated dataset contains reads from a mixture of 1,000 reference haplotypes (each an operational taxonomic unit: OTU). The frequencies at which reads were generated from contributing references were varied to create datasets with a range of Shannon Diversity. As diversity increases the frequency distribution begins to approach a uniform distribution.

**Figure S6:**
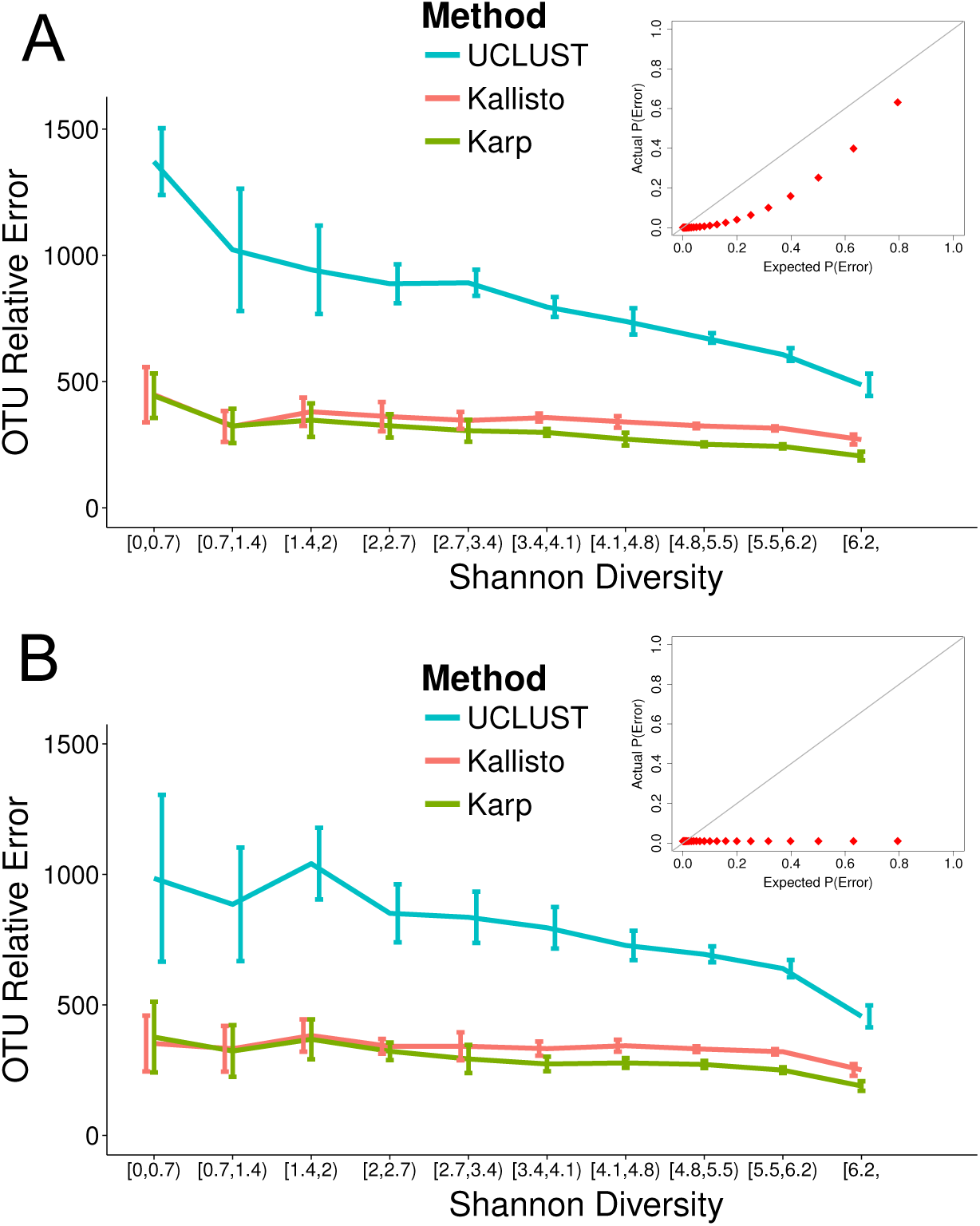
Impact of assumption that base-quality scores accurately represent probability of sequencing error. For two different models of sequencing error we simulated 50 samples and classified them with Karp, Kallisto, and UCLUST/USEARCH. Each method is represented by a different colored line, and bars represent 95% confidence intervals (A) In our first model the true rate of sequencing error varied with the base-quality score, but was smaller than Karp’s model assumes. (B) In our second model, errors were distributed uniformly at 1% of bases in each read, independent of whatever base-quality score was assigned.

**Figure S7:**
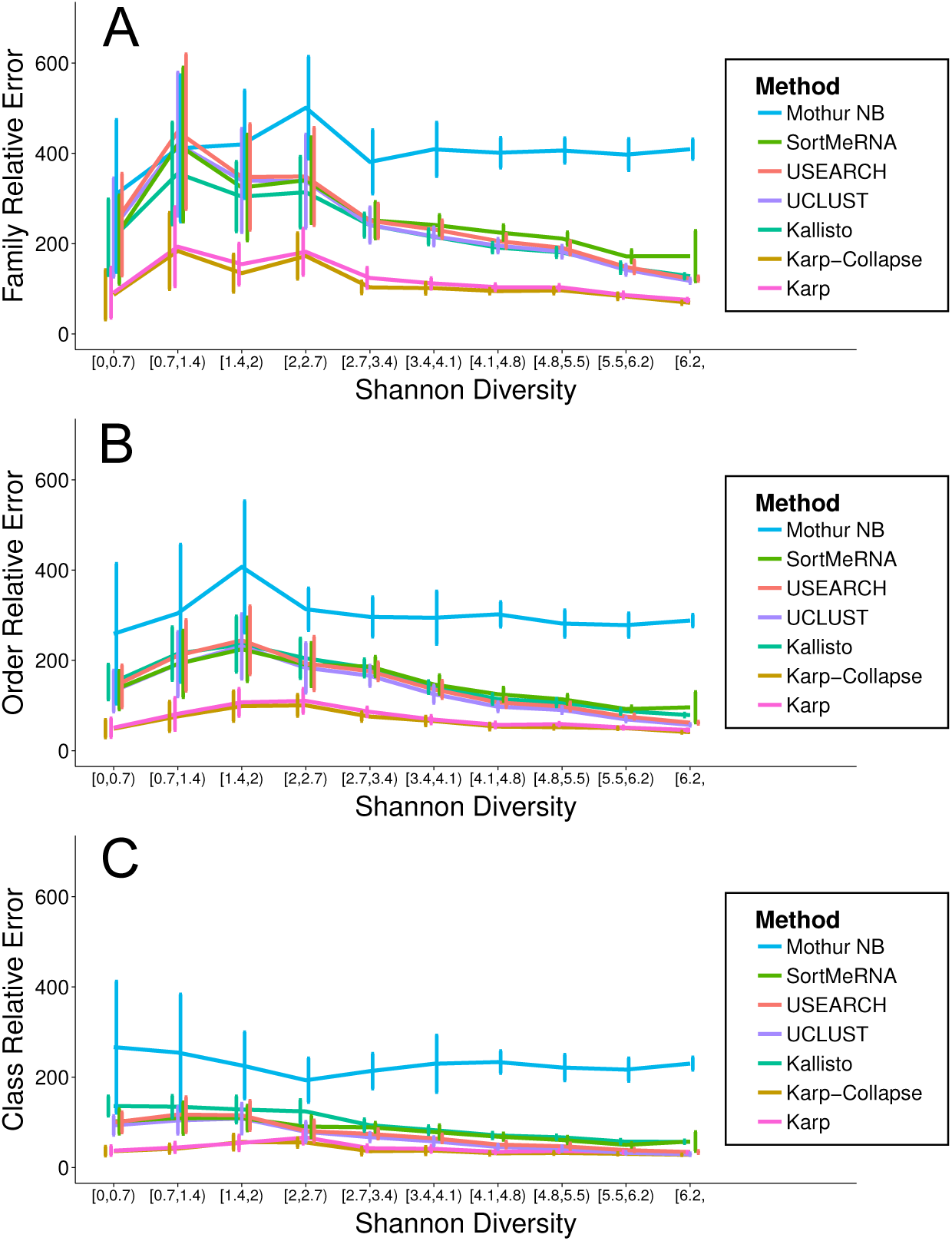
Average absolute error and 95% confidence intervals from the taxonomic classification of 110 simulated samples with 75bp paired-end reads. Taxonomy was classified using Karp, Kallisto, UCLUST, USEARCH, SortMeRNA, and the Naive Bayes method implemented in Mothur. Counts were aggregated for OTUs classified in the same (A) Family, (B) Order, or (C) Class and taxa with a frequency > 0.1% were compared to their true counts.

**Figure S8:**
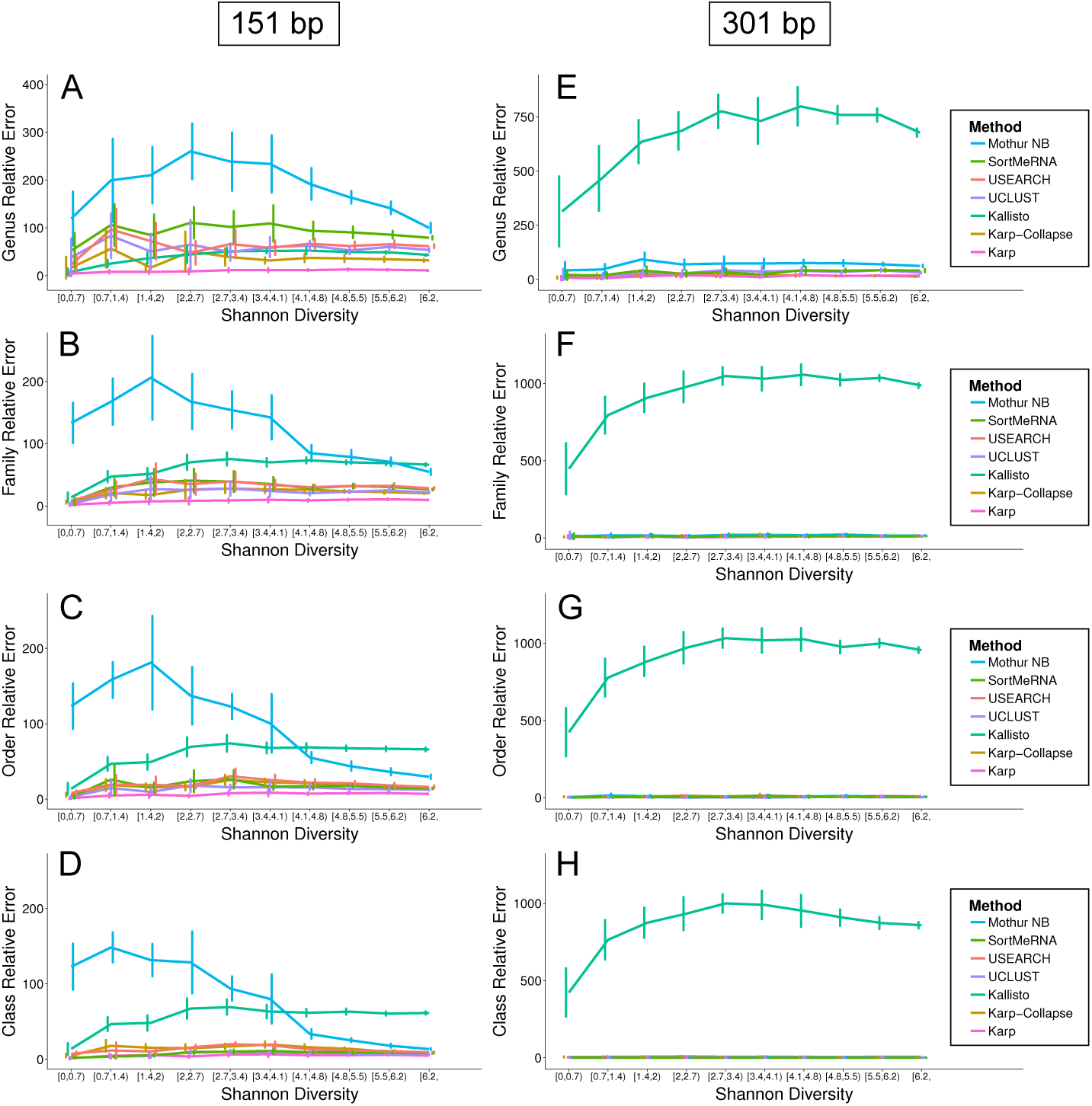
Average absolute error and 95% confidence intervals from the taxonomic classification of simulated paired-end read samples. Taxonomy was classified using Karp, Kallisto, UCLUST, USEARCH, SortMeRNA, and for the 151bp samples the Naive Bayes method implemented in Mothur. For 151bp paired-end reads counts were aggregated for OTUs classified in the same (A) Genus, (B) Family, (C) Order, or (D) Class and taxa with a frequency > 0.1% were compared to their true counts. Likewise, for 301bp paired-end reads counts were aggregated for OTUs classified in the same (E) Genus, (F) Family, (G) Order, or (H) Class and taxa with a frequency > 0.1% were compared to their true counts.

